# An iPSC-based experimental framework for investigating phenotypic variation associated with Jomon ancestry in the present-day Japanese population

**DOI:** 10.64898/2026.10.07.756619

**Authors:** Yuka Nakamura, Yusuke Watanabe, Hideyuki Tanabe, Asei Hirai, Koichi Higashi, Ken Kurokawa, Atsushi Toyoda, Tomoaki Kato, Masayoshi Tsukahara, Kae Koganebuchi, Yusuke Hamazaki, Takanori Takebe, Masanori Imamura, Takafumi Ishida, Jun Ohashi, Hiroki Oota

**Author notes:** These authors contributed equally to this work (Yuka Nakamura, Yusuke Watanabe).

## Abstract

Ancestry-related genomic variation may contribute to phenotypic diversity. However, its cellular consequences remain difficult to evaluate. To address this, we developed a human induced pluripotent stem cell (hiPSC)-based framework for investigating such phenotypes. We used the Japanese population as a model of Jomon-related and continental East Asian ancestry admixture. Jomon ancestry, estimated using the Jomon allele score (JAS), was associated with clinical measures, including renal-related traits and serum triglycerides in 34,152 Tohoku Medical Megabank participants. Peripheral blood mononuclear cell transcriptomics of donors from the upper and lower extremes of the JAS distribution (n = 10) suggested group-level differences in immune-related gene expression. To test whether the serum triglyceride association coincided with hepatic cellular phenotypes, we differentiated hiPSCs from these donors into hepatocyte-like cells. Neither intracellular triglycerides nor lipid-related gene expression differed between groups. These findings highlight the value of a genetically characterized hiPSC-based framework for investigating ancestry-associated phenotypes across cellular contexts.

## Introduction

Genome-wide association studies (GWAS) have identified numerous genetic variants associated with clinical phenotypes^1-4^, including variants whose allele frequencies differ among populations. Such associations can contribute to the development of precision medicine and provide insight into the evolutionary history of human phenotypic diversity^5-7^. For many associated loci, however, the causal variants, target genes, relevant cell types, and cellular mechanisms remain unresolved^8,9^. Animal models are indispensable for functional studies but may not reproduce effects that depend on human-specific regulatory architecture or broader ancestry-related genomic context^10^. Human induced pluripotent stem cells (hiPSCs) provide a complementary experimental system because they retain donor-specific genomic backgrounds and can be differentiated into multiple cell types under controlled conditions^11-15^. Panels of genetically characterized hiPSC lines may therefore enable systematic investigation of cellular phenotypes associated with ancestry-related variation.

Genomic variation in present-day Japanese populations primarily reflects contributions from Jomon-related and continental East Asian ancestries^16,17^. This dual-structure model was first proposed based on morphological studies and has since been strongly supported by genetic evidence^18-28^. The ancestral lineages contributing to these components experienced long periods of separation before their admixture in the Japanese archipelago. They also differed in demographic history and subsistence, including foraging and agricultural lifeways^29,30^. These differences may have contributed to variation in allele frequencies at loci involved in metabolism, immunity, and other physiological processes^31,32^. Subsequent admixture and demographic history resulted in interindividual and regional variation in Jomon-related ancestry across the Japanese archipelago, providing an opportunity to examine associations between ancestry and phenotypic variation within the present-day population^16,17,33,34^. Previous studies have associated higher estimated Jomon ancestry with body mass index (BMI) and have proposed Jomon-enriched haplotypes related to triglyceride metabolism^16,33^. These findings raise the possibility that differences in Jomon-related ancestry may be associated with phenotypic variation among present-day Japanese individuals.

However, population-level associations do not establish the underlying cellular mechanisms, and functional studies of Jomon-related genomic variation remain limited. The Jomon allele score (JAS)^16^, a genome-wide summary score based on alleles inferred to be Jomon-derived, provides a quantitative proxy for comparing Jomon-related ancestry among individuals. Combining this score with donor-derived cellular models could therefore provide a tractable framework for exploring whether ancestry-associated clinical traits are accompanied by measurable cellular phenotypes.

In this study, we aimed to establish an initial hiPSC-based framework for evaluating the phenotypic diversity associated with Jomon ancestry. We first tested associations between JAS and clinical measures using genomic and clinical data from the Tohoku Medical Megabank Organization (ToMMo)^35,36^. For renal-related traits, we intersected trait-associated GWAS loci with previously reported XP-nSL signals contrasting Jomon and Han Chinese in Beijing (CHB) populations to nominate candidate regions for further investigation^37^. We then selected donors from the upper and lower extremes of the JAS distribution, characterized basal gene expression in their peripheral blood mononuclear cells (PBMCs), and established hiPSC lines from the same donors. As an initial application of this resource, we focused on the observed association between JAS and serum triglyceride levels. We differentiated the hiPSCs into hepatocyte-like cells (HLCs) and assessed intracellular triglyceride content and the expression of *ACACA, APOB, CPT1A, FASN,* and *PDK4*.

Together, these analyses establish an initial experimental framework and a JAS-stratified hiPSC panel for investigating cellular phenotypes associated with ancestry-related genomic variation.

## Results

### Jomon ancestry is associated with diverse clinical traits, including renal-related traits and serum triglyceride levels

To quantify interindividual variation in Jomon ancestry, we calculated the Jomon allele score (JAS) for 60,803 individuals using genomic data of ToMMo. JAS showed broad interindividual variation (Figure 1A and 1B), consistent with the distribution reported for a commercial genome database in the previous study^16^. This broad distribution allowed us to systematically investigate associations between Jomon ancestry and phenotypic variation.

**Figure 1:**
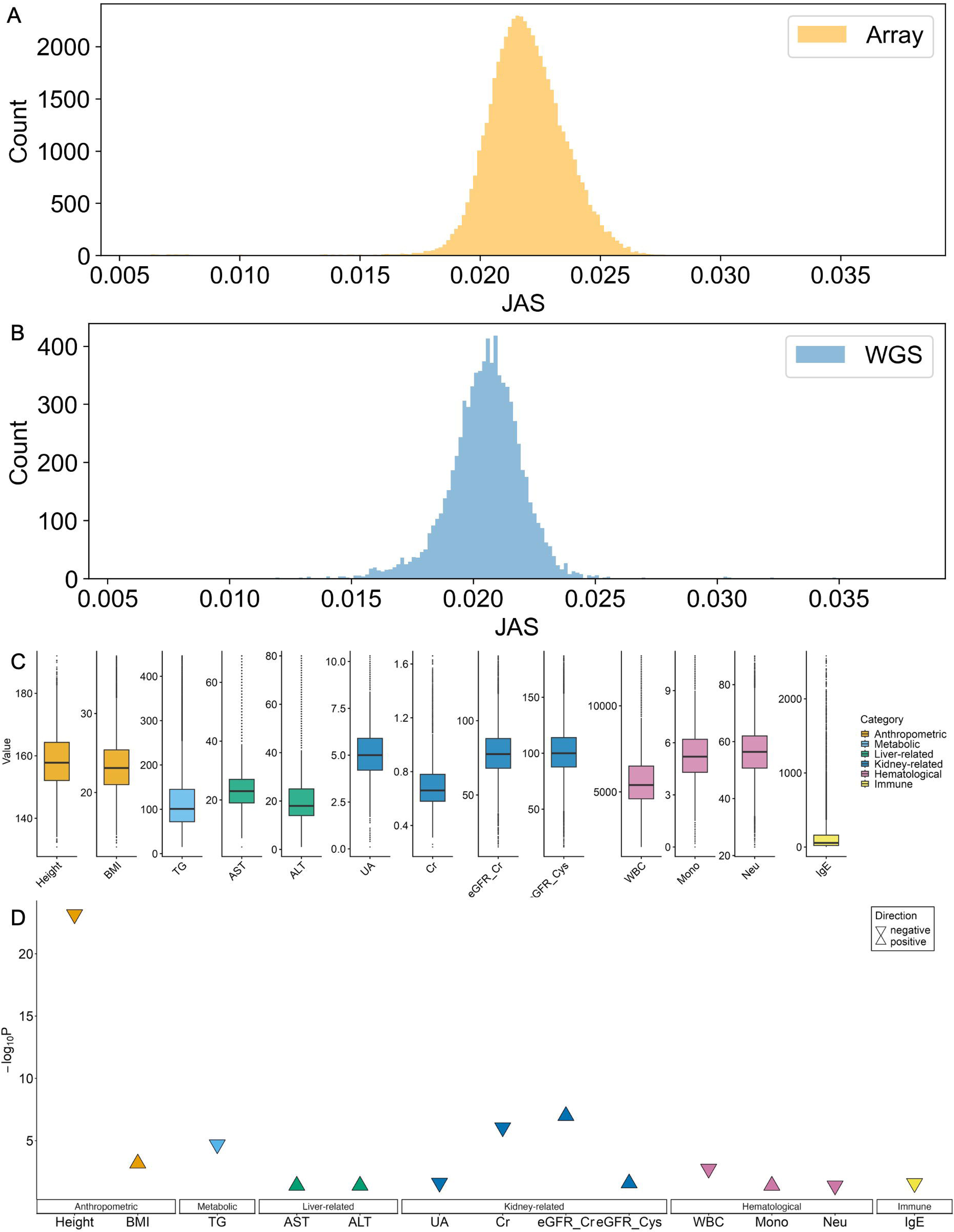
Jomon ancestry is associated with clinical phenotypes in a large Japanese cohort. (A) Distribution of JAS in the cohort with SNP array data. (B) Distribution of JAS in the cohort with WGS data. (C) Distribution of clinical values. Data points exceeding four standard deviations (4SD) from the mean were excluded prior to plotting. (D) Association between JAS and clinical parameters analyzed in this study. Linear regression analyses were performed between JAS and 31 clinical parameters in 34,152 participants, of which 13 remained significant after Bonferroni correction (adjusted p-value < 0.05). The upward- and downward-pointing triangles indicate positive and negative associations, respectively. The y-axis shows the statistical significance of each association (−log10(p-value)). Results for all analyzed parameters are shown in Figures S1A and S1B.

Next, we evaluated associations between JAS and clinical test values using linear regression analyses (Tables S1 and S2). Of the 31 parameters tested, 13 remained significantly associated with JAS after Bonferroni correction (adjusted p-value < 0.05). Specifically, higher JAS was associated with higher values for BMI, eGFR based on serum creatinine (eGFRcr), eGFR based on serum cystatin C (eGFRcys), AST, ALT, and monocytes, and with lower values of height, uric acid (UA), creatinine, triglycerides (TG), white blood cell count, neutrophils, and IgE (Table 1, Figures 1C, 1D, S1A, S1B, and S1C).

**Table 1.** Results of regression analyses of Jomon ancestry and clinical traits.

| Trait | Unit | Estimate | P_value | P_adj |
| --- | --- | --- | --- | --- |
| Height | cm | -198.71 | 2.23E-25 | 6.93E-24 |
| Weight | kg | -39.30 | 0.20801 | 0.29311 |
| Weight (Age20) | kg | -2.32 | 0.92352 | 0.92352 |
| Waist | cm | 27.69 | 0.35858 | 0.46316 |
| BMI | kg/m2 | 44.22 | 0.00011 | 0.00067 |
| UN | mg/dL | 17.24 | 0.17481 | 0.25806 |
| UA | mg/dL | -10.98 | 0.00552 | 0.02446 |
| Cr | mg/dL | -2.30 | 8.41E-08 | 8.70E-07 |
| eGFRcr | ml/min/1.73m2 | 292.74 | 6.60E-09 | 1.02E-07 |
| eGFRcys | ml/min/1.73m2 | 156.52 | 0.00756 | 0.02604 |
| Glu | mg/dL | 122.50 | 0.03709 | 0.07186 |
| HDL | mg/dL | 89.59 | 0.09801 | 0.15991 |
| LDL | mg/dL | -72.47 | 0.48198 | 0.57466 |
| TG | mg/dL | -1068.44 | 2.61E-06 | 2.02E-05 |
| γ-GTP | U/L | 138.00 | 0.13519 | 0.20954 |
| AST | U/L | 60.96 | 0.01659 | 0.04047 |
| ALT | U/L | 90.02 | 0.01194 | 0.03702 |
| SBP | mmHg | 125.17 | 0.02602 | 0.05377 |
| DBP | mmHg | 40.76 | 0.23686 | 0.31924 |
| RBC | 10,000 / μL | 110.25 | 0.38456 | 0.47685 |
| Hb | g/dL | -1.03 | 0.78214 | 0.81903 |
| Ht | % | 3.27 | 0.75649 | 0.81903 |
| WBC | /μL | -17891.44 | 0.00035 | 0.00182 |
| HbA1c | % | -0.37 | 0.79261 | 0.81903 |
| PLT | 10,000 / μL | -9.45 | 0.59611 | 0.68442 |
| Lym | % | 48.69 | 0.07684 | 0.13234 |
| Mono | % | 10.90 | 0.01533 | 0.04047 |
| Eos | % | 13.56 | 0.02489 | 0.05377 |
| Bas | % | 1.94 | 0.05403 | 0.09852 |
| Neu | % | -70.36 | 0.01697 | 0.04047 |
| IgE | IU/mL | -2707.00 | 0.00707 | 0.02604 |

Notably, we found associations with renal-related traits that had not been highlighted in previous studies of Jomon-related ancestry^16,17,33,34^. Higher JAS was associated with lower levels of serum creatinine and UA, as well as higher eGFRcr and eGFRcys. Because eGFRcr was derived from serum creatinine, the associations with serum creatinine and eGFRcr were not independent.

We examined the overlap between renal-related GWAS signals and genomic regions with extreme XP-nSL signals and prioritized five candidate loci: *ABCG2*, the *MHC* region, *MYL2*, *VHRT*, and *WDR1* (Tables S3 and S4). Among these loci, we prioritized *ABCG2* due to its established role in urate transport^38-40^. For the seven variants with the largest allele frequency differences between Jomon and CHB in the *ABCG2* region, alleles enriched in Jomon individuals were consistently associated with lower UA levels (Table S5), consistent with the inverse association between JAS and serum UA (Figures 1C and D). The G allele of rs2231157, an intronic variant in *ABCG2,* was more frequent in Jomon individuals than in worldwide populations examined (Table S6). Regulatory annotation indicated that rs2231157 overlapped regulatory features potentially related to *ABCG2* expression (Table S7). These analyses were intended to prioritize candidate variants for further investigation rather than to identify causal variants.

In addition to renal traits, our analysis revealed a significant association between Jomon ancestry and serum TG levels. The association with TG was opposite in direction to that reported in a previous study^16^; higher JAS was associated with lower TG levels. We therefore selected serum TG as an initial test case for the hiPSC resource because lipid-related cellular phenotypes, particularly intracellular TG accumulation, can be assessed in hiPSC-derived hepatocyte-like cells.

### PBMC transcriptomes suggest group-level differences in immune-related gene expression between upper- and lower-JAS groups

To explore whether donors with contrasting JAS values also showed differences in PBMC transcriptional profiles, we performed an exploratory transcriptomic analysis. To maximize the contrast in JAS, we selected 10 individuals representing the two extremes of the JAS distribution, regardless of age or sex (Table S8, S9, Figure 2A, and 2B). The "upper-JAS group" consisted of the five individuals selected from the upper extreme of the JAS distribution (JAS range: 0.0302–0.0378), while the "lower-JAS group" included the five individuals selected from the lower extreme of the JAS distribution (JAS range: 0.0058–0.0061). Individual JAS values are provided in Table S8 and S9.

**Figure 2:**
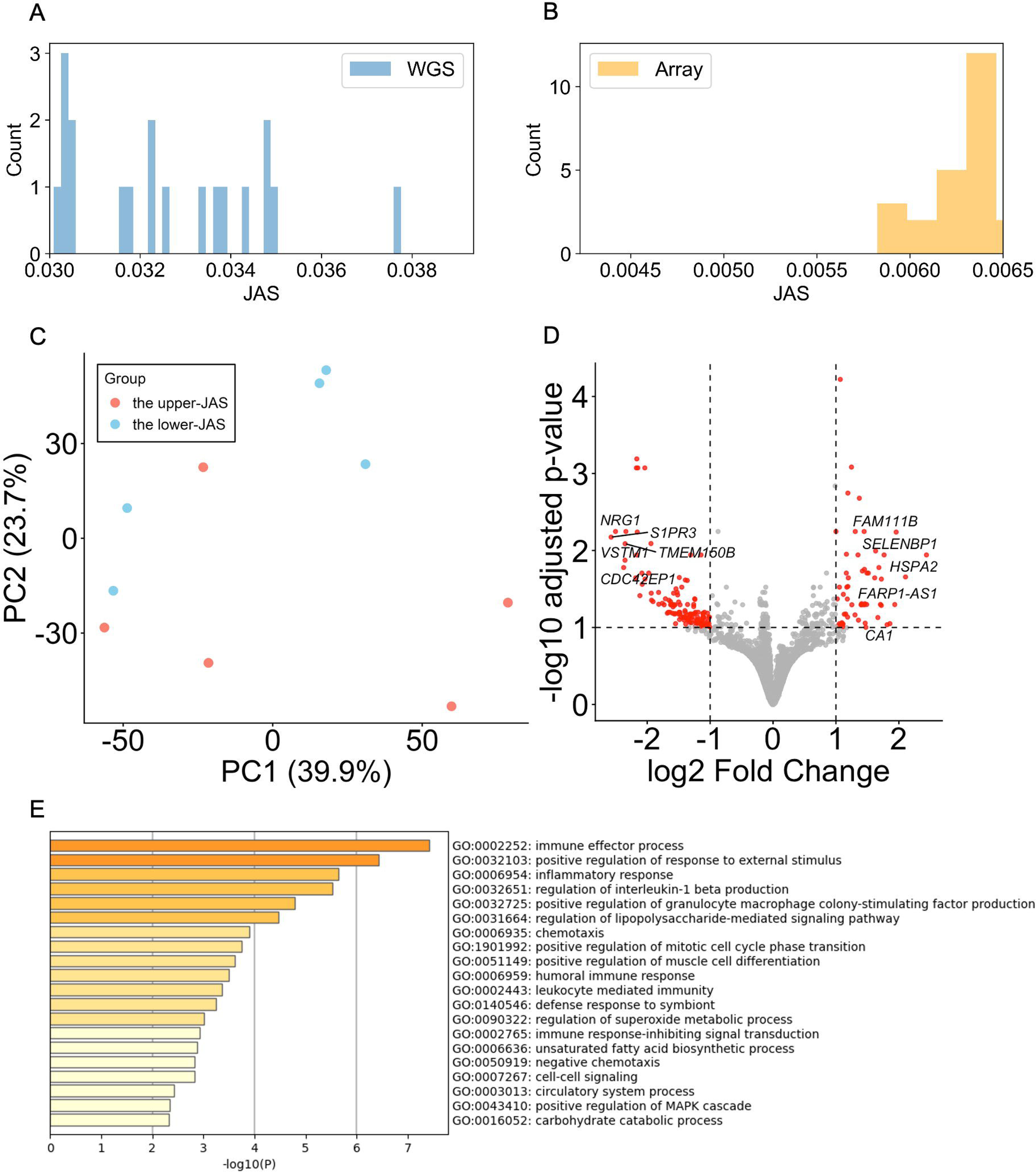
PBMC transcriptomes reveal differences in gene expression between the upper-JAS (n = 5) and lower-JAS (n = 5) groups. (A) Enlarged view of the lower range of the JAS distribution shown in Figure 1A. (B) Enlarged view of the higher range of the JAS distribution shown in Figure 1B. (C) PCA of gene expression for 10 individuals. Red dots represent the upper-JAS group. Blue dots represent the lower-JAS group. (D) Volcano plot of DEGs between the upper- and lower-JAS groups. Red dots indicate significantly differentially expressed genes (DEGs) with an adjusted p-value < 0.1 and an absolute log2 fold change (|log2FC|) ≥ 1. The top 5 up-regulated and down-regulated genes are labeled with their symbols. (E) GO Biological Process terms enriched among genes downregulated in the lower-JAS group relative to the upper-JAS group.

To characterize the molecular state of the donor cells used in this study, we compared the gene expression profiles of PBMCs between the upper-JAS (n = 5) and lower-JAS (n = 5) groups after quality control (Table S10). Principal component analysis (PCA) showed no clear separation between the groups (Figure 2C). Differentially expressed genes (DEGs) were defined using an adjusted p-value < 0.1 and |log2FC| ≥ 1. A total of 164 genes met these criteria, with 51 showing higher and 113 showing lower expression in the lower-JAS group relative to the upper-JAS group (Tables S11, S12, and Figure 2D). Functional enrichment analysis was performed to characterize these DEGs, using the 14,562 filtered gene entries as the background set. No significant enrichment was detected among genes with higher expression in the lower-JAS group (Table S13). In contrast, genes with lower expression were significantly enriched for immune-related biological processes, including inflammatory response and positive regulation of response to external stimulus (Table S14 and Figure 2E). These exploratory results suggest group-level differences in the expression of immune-related genes.

Given the association between JAS and monocyte proportions observed in clinical analyses (Table 1), we assessed whether differences in immune cell composition could explain the observed DEGs. CIBERSORT analysis did not detect significant differences in estimated immune cell proportions between the upper- and lower-JAS groups for evaluable cell types, either before or after Benjamini–Hochberg correction (p > 0.05; Table S15 and Figure S2)^41^, although effects of cell composition cannot be excluded. Given the small sample size (n = 10) and unmatched age and sex distribution, these findings should be interpreted as exploratory group-level differences.

### hiPSC lines from upper- and lower-JAS donors retain pluripotency

To establish a cellular resource for investigating the phenotypic variation associated with Jomon ancestry, we generated hiPSCs from PBMCs of individuals in the upper-JAS and lower-JAS groups. We established 10 hiPSC lines, five from each group, and characterized them to assess pluripotent marker expression and differentiation potential. All established hiPSC lines expressed key pluripotency markers (Figures 3A, 3B, and S3). Direct differentiation into the three germ layers demonstrated the expression of ectodermal, mesodermal, and endodermal markers, respectively (Figures 3C and S4–6). Differentiation potential was also demonstrated by the generation of cardiomyocyte-like cells (Figures 3D and S7). To assess clearance of the Sendai virus (SeV) vector used for reprogramming, we performed qPCR for SeV vector sequences. While one line (T2) showed residual vector presence at passage 4 (P4), it became negative by P10. All other lines were negative for the vector at an early stage (Table S16). Cytogenetic analysis using QFH-banding and multicolor fluorescence in situ hybridization (mFISH) revealed no numerical or structural chromosomal abnormalities (Figure 3E and S8).

**Figure 3:**
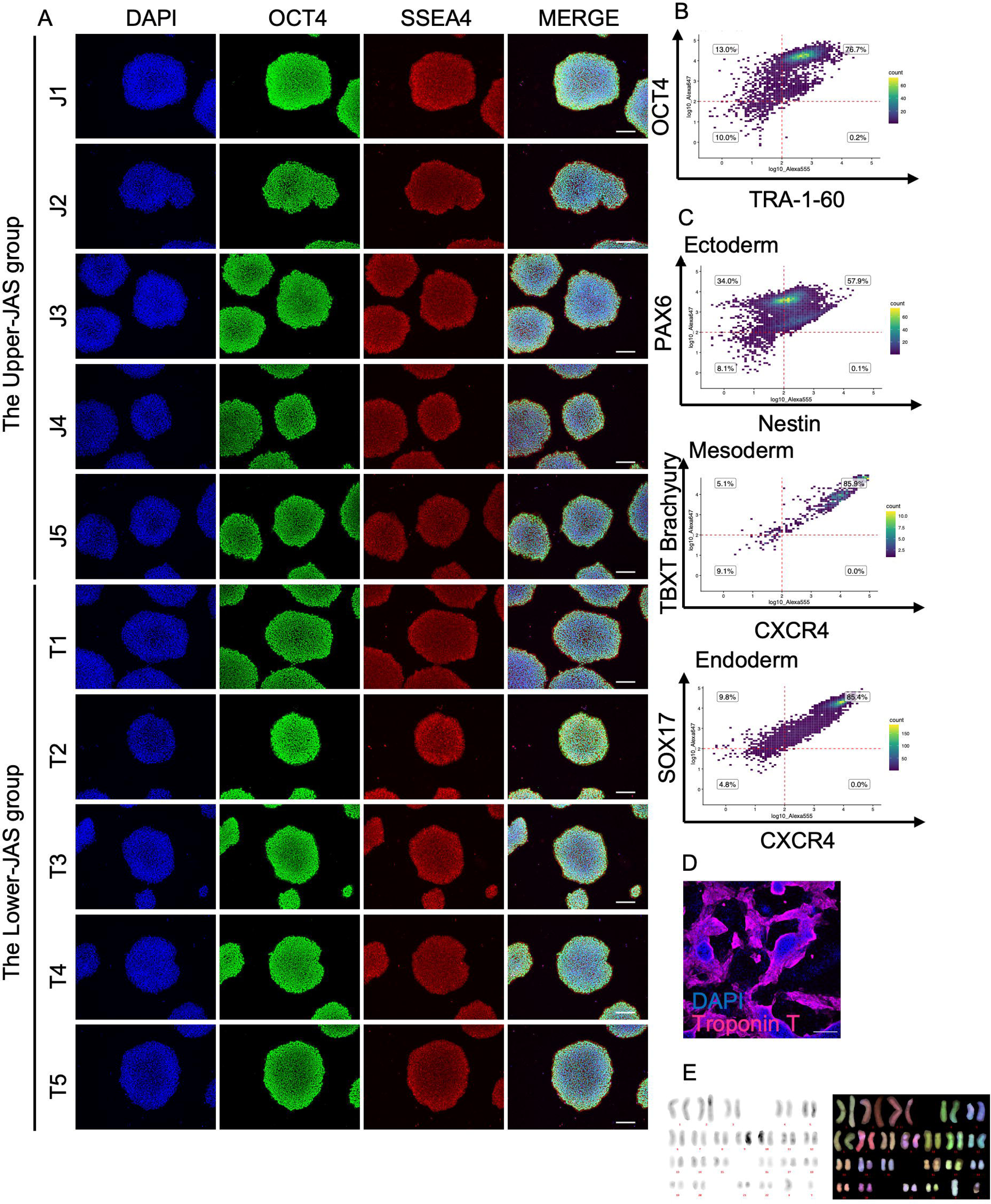
Characterization and differentiation potential of hiPSC lines from PBMCs of donors in the upper- and lower-JAS groups. (A) Immunofluorescence images of pluripotency markers (OCT4 and SSEA4) in hiPSCs. Scale bar: 200 μm. (B) Representative flow cytometric analysis of pluripotency markers (OCT4 and TRA-1-60). (C) Representative flow cytometric analysis of ectodermal (PAX6 and Nestin), mesodermal (TBXT (Brachyury) and CXCR4), and endodermal markers (SOX17 and CXCR4). (D) Immunofluorescence images of the cardiomyocyte marker (Troponin T) in hiPSC-derived cardiomyocytes. Scale bar: 200 μm. (E) Representative QFH-banding and mFISH images of chromosomes from metaphase spreads of hiPSCs.

To verify sample identity, we compared genotypes at 10 short tandem repeat (STR) loci between the iPSC lines and their parental PBMCs. The STR profiles of the hiPSCs matched those of the parental PBMCs in all evaluable PBMC–hiPSC pairs (Table S17), although a parental PBMC profile was unavailable for line J2. Finally, all hiPSC lines tested negative for mycoplasma by the PCR-based assays (Table S18). Together, these quality-control assessments indicated that the hiPSC lines were suitable for downstream differentiation and cellular phenotyping.

### hiPSC-derived HLCs from upper- and lower-JAS donors show no detectable differences in intracellular TG content or *ACACA, APOB, CPT1A, FASN,* and *PDK4* expression

To examine whether the association between JAS and serum TG levels was accompanied by differences in hepatic cellular phenotypes, we differentiated the hiPSCs into hepatocyte-like cells (HLCs) (Figure 4A). Differentiation toward HLCs was assessed by immunocytochemistry and qPCR (Table S19). Immunocytochemistry showed expression of the hepatocyte markers albumin and HNF4α (Figure 4B and S9). The mean HNF4α-positive cell rate across the 10 lines was 62.5 ± 18.3%, ranging from 26.8% to 85.9% (Table S20 and Figure 4C). No significant difference in HNF4α-positive rate was observed between the upper- and lower-JAS groups. qPCR analysis showed increased expression of hepatocyte markers after differentiation (Figure 4D). Together, these results supported hepatocyte-like differentiation across the JAS-stratified hiPSC panel.

**Figure 4:**
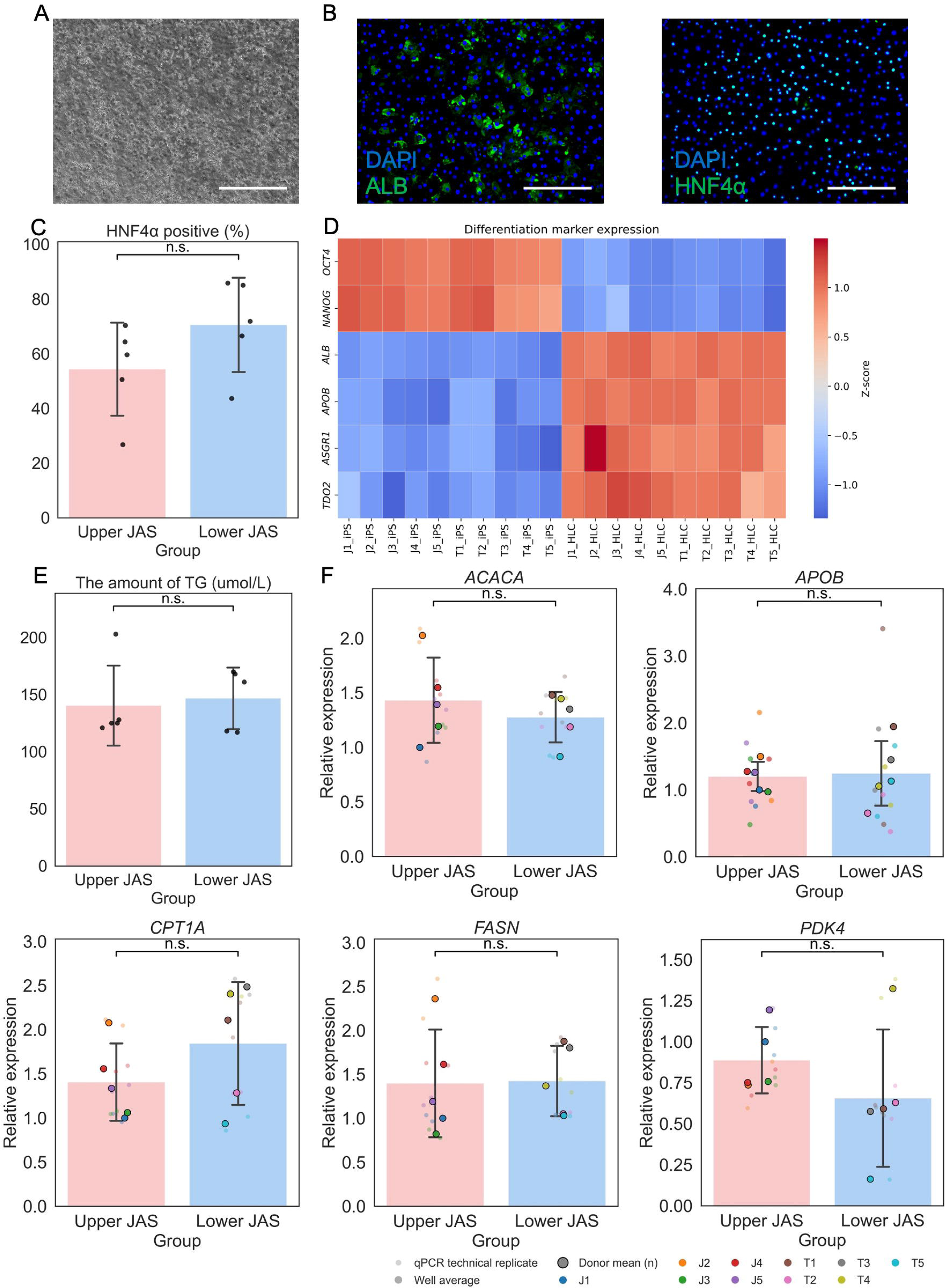
Functional evaluation of triglyceride-associated phenotypes in hiPSC-derived hepatocyte-like cells (HLCs). (A) Representative phase-contrast images of hiPSC-derived HLCs. Scale bars: 200 μm. (B) Representative immunofluorescence images of HLCs showing the expression of hepatic markers albumin (ALB) and HNF4α. Scale bars: 200 μm. (C) Quantification of HNF4α-positive cells across all HLC lines. Each dot represents an independent hiPSC line. No significant difference was observed between the upper- and lower-JAS groups by the Mann–Whitney U test (n = 5 independent hiPSC lines per group). (D) Heatmap showing Z-score-normalized expression levels of hepatic marker genes before and after hepatic differentiation, measured by qPCR (n = 5 independent hiPSC lines per group). (E) Intracellular triglyceride content in HLCs derived from individuals with the upper- and lower-JAS groups. No significant difference was observed between the upper- and lower-JAS groups by the Mann–Whitney U test (n = 5 independent hiPSC lines per group, with four wells pooled for each line). (F) The expression levels of *ACACA, APOB, CPT1A, FASN,* and *PDK4* in HLCs derived from individuals with the upper- and lower-JAS groups are shown relative to the upper-JAS group (set to 1). Large dots indicate the mean value for each independent hiPSC line. For *APOB*, medium-sized dots indicate individual well-level measurements, whereas for the other genes, small dots indicate individual qPCR replicate measurements. No significant differences were observed between the upper- and lower-JAS groups by the Mann–Whitney U test (n = 5 independent hiPSC lines per group).

To investigate whether the association between JAS and serum TG levels was accompanied by differences in hepatic lipid-related phenotypes, we measured intracellular TG content and the expression of *ACACA, APOB, CPT1A, FASN,* and *PDK4* as lipid-related cellular readouts. No significant differences were observed in intracellular TG content or the expression of these five genes between the upper- and lower-JAS groups (Table S20, S21, Figures 4E, 4F, and S10). Under the present culture conditions, we found no evidence of differences between the upper- and lower-JAS groups in the hepatic phenotypes examined.

## Discussion

In this study, we aimed to establish an initial hiPSC-based framework for evaluating the phenotypic diversity associated with Jomon ancestry in the present-day Japanese population. To achieve this, we integrated large-scale association analyses of clinical measures, transcriptomic profiling of PBMCs, and the generation of the JAS-stratified hiPSC panel for functional evaluation. Using JAS as a quantitative proxy for Jomon-related ancestry, we identified associations with multiple clinical phenotypes and established an initial experimental framework for investigating ancestry-associated cellular phenotypes. The JAS-stratified hiPSC panel established in this study provides a human cellular model incorporating interindividual variation in JAS among present-day Japanese population.

Our regression analysis showed significant associations between JAS and several renal-related traits that have not been highlighted in previous studies of Jomon-related ancestry. This finding represents a previously undocumented link between Jomon ancestry and modern physiological traits. Specifically, higher JAS was associated with lower levels of serum uric acid and creatinine and higher eGFR (eGFRcys and eGFRcr). While UA acts as a potent antioxidant, its excessive accumulation can lead to renal toxicity^42^. Our analyses prioritize candidate loci rather than establish causal or adaptive variants, and functional studies will be required to determine their regulatory effects. The JAS-stratified hiPSC panel established here could support future functional studies of candidate loci related to urate metabolism in relevant differentiated cell types.

CIBERSORT-based estimation did not detect significant differences in immune cell composition between the upper- and lower-JAS groups, despite the association between JAS and monocyte proportions in the large clinical cohort. The difference between these observations may reflect the much smaller sample size and limited statistical power of the PBMC analysis. The exploratory PBMC comparison suggests lower expression of a subset of immune-related genes in the lower-JAS group. These exploratory expression patterns are not directly comparable with previous genetic predictions of immune-related adaptation because they represent transcriptional states in unstimulated PBMCs^16^. Given the small sample size and differences in age and sex between the groups, these differences cannot be attributed specifically to JAS. These transcriptomic trends provide a basis for future studies using larger, better-matched cohorts.

We also observed associations between JAS and anthropometric and metabolic traits. The observed positive association between JAS and BMI aligns with previous in silico predictions^16,33,34^. The inverse association between JAS and height is also compatible with morphological reports indicating shorter stature among Jomon individuals than among continental migrants^43,44^.

Intriguingly, although BMI and TG are typically positively correlated in general populations^45,46^, they exhibited opposite associations with JAS in our study. The inverse association between JAS and serum TG contrasts with ancestry-based predictions in present-day Japanese populations^16^ and with the recent ancient-genome analysis by Watanabe et al.^37^, which identified Jomon-lineage selection signals at loci related to body size and lipid metabolism, including *FTO* and *ZPR1–APOA5*. In that study, the Jomon-selected haplotypes at these loci were associated with increased BMI and TG, respectively. Thus, the agreement for BMI but not TG suggests that a genome-wide ancestry score in an admixed present-day population does not necessarily recapitulate locus-specific effects inferred from ancient Jomon genomes^37^. Because BMI incorporates height, its positive association with JAS may also partly reflect the inverse association between JAS and height rather than lipid metabolism alone. The association between JAS and serum TG levels motivated us to test whether corresponding hepatic cellular phenotypes could be detected using hiPSC-derived HLCs.

To explore this association at the cellular level, we used our newly established JAS-stratified hiPSC resource, comprising five lines each from the upper- and lower-JAS groups. As noted previously, our regression analysis revealed that higher JAS was associated with lower levels of serum TG. Serum TG levels reflect multiple processes, including hepatic lipid production and lipoprotein secretion. To test this hypothesis at the cellular level, we focused on the hepatic contribution to TG metabolism and differentiated the hiPSCs into HLCs. We compared intracellular TG content and the expression of *ACACA, APOB, CPT1A, FASN,* and *PDK4*. However, no significant differences were observed between the upper- and lower-JAS groups for either of these readouts. Because this study did not assess TG secretion or the secretion of lipid-related proteins, and the maturation state of hiPSC-derived HLCs remains lower than that of adult hepatocytes, subtle differences associated with JAS may not be fully captured. Thus, the association between JAS and serum TG levels was not recapitulated by the hepatic phenotypes examined here and may involve systemic regulation or other hepatic processes not captured by these assays. Together, these findings underscore the complexity of the relationship between ancestry, clinical phenotypes, and cellular phenotypes, highlighting the need for a framework that leverages multiple genetically characterized hiPSC lines to investigate ancestry-associated phenotypes across diverse cellular contexts. This framework can be extended to larger numbers of genetically characterized hiPSC lines and diverse differentiated cell types to determine whether additional ancestry-associated phenotypes emerge and to clarify the mechanisms underlying association between JAS and lipid-related phenotypes.

In conclusion, moving beyond population-level association analyses, this study establishes an initial experimental framework using donor-derived hiPSCs to investigate cellular phenotypes associated with genome-wide ancestry-related genomic variation rather than individual loci. Our findings further suggest that population-level associations may not be readily recapitulated as detectable cellular phenotypes under a single experimental condition, motivating expansion of the resource to a larger panel of genetically characterized donors and diverse differentiated cell types. Such an expanded framework could provide a foundation for investigating how ancestry-related genomic diversity contributes to phenotypic variation at the cellular level.

### Limitations of the study

First, the exploratory PBMC RNA-seq analysis included only five donors in each JAS group, and the groups were not matched for age or sex. Therefore, the observed differences in immune-related gene expression cannot be attributed specifically to Jomon ancestry. Second, the number of hiPSC lines used for functional analyses was limited to five lines per JAS group. Because donor-to-donor genetic variation may influence cellular phenotypes independently of Jomon ancestry, larger panels of genetically characterized hiPSC lines will be needed to distinguish ancestry-associated effects from donor-specific variation. Finally, our assessment of hepatic lipid phenotypes was limited to intracellular TG accumulation and the expression of *ACACA, APOB, CPT1A, FASN,* and *PDK4*, and we did not assess TG secretion or the secretion of lipid-related proteins. In addition, hiPSC-derived HLCs remain less mature than adult hepatocytes, which may have limited our ability to detect subtle ancestry-associated phenotypes. Future studies using more mature hepatocyte models and incorporating additional measures of hepatic lipid metabolism may enable a more comprehensive assessment of ancestry-associated hepatic phenotypes.

## Materials and Methods

### Experimental model and study participant details Ethics statement

Genome data and clinical tests were provided by the Tohoku Medical Megabank Organization^35,36,47^. All donor samples were obtained from the ToMMo Biobank. The generation and use of hiPSCs were approved by The University of Tokyo (approved No. 24-489) and ToMMo (approved No. 2021-0062-1). The experiment involving genetic modification was approved by Kyoto University (approved No. 240444).

### Cell lines

The 10 hiPSC lines used in this study were established from PBMCs of healthy Japanese donors provided by the ToMMo. PBMC samples separated by density gradient centrifugation were obtained from ToMMo. These donors were selected based on their JAS, a genome-wide quantitative proxy for Jomon-related Jomon ancestry. The upper-JAS group," consisting of the five individuals, and the "lower-JAS group comprised the five donors with JAS values of 0.0302–0.0378, whereas the lower-JAS group comprised five donors with JAS values of 0.0058–0.0061. The hiPSC lines derived from these individuals were designated J1–J5 and T1–T5, respectively. To ensure a high-resolution genomic background for downstream functional analyses, donors for the upper-JAS group were selected exclusively from a WGS dataset comprising 7,908 samples. In contrast, the lower-JAS group was selected from a combined pool of a genome-wide imputation dataset of 52,895 samples to prioritize the inclusion of individuals with the most extreme low-ancestry scores.

## Method details

### Calculating the Jomon allele score

To identify individuals with high and low proportions of Jomon genetic ancestry, we utilized the JAS as a quantitative metric. The JAS for each sample was calculated using genomic data as previously described^16^. Briefly, this method estimates the ancestral component by counting Jomon-specific derived alleles across the genome. In this study, JAS was computed for two distinct datasets from ToMMo: (1) a whole-genome sequencing (WGS) dataset consisting of 7,908 samples^48^, and (2) a genome-wide imputation dataset of 52,895 samples^47,49^. For the WGS dataset, 198,244 SNPs reported by Watanabe and Ohashi (2023)^16^ were used for JAS calculation, while 183,323 SNPs from the same report were applied to the imputation dataset.

### Regression analysis of clinical tests and JAS

To identify phenotypes associated with Jomon ancestry, linear regression analyses were conducted between clinical test values and JAS. Out of 52,895 samples, several exclusion criteria were applied to ensure data quality and homogeneity: (1) individuals estimated to be within second-degree kinship using PLINK 2.0^50,51^, (2) individuals with reported non-East Asian ancestry based on self-reported questionnaires, and (3) individuals with missing data among the variables included in the analysis. Consequently, 34,152 samples were included in the final analysis. Data points deviating by more than four standard deviations (4SD) from the mean were identified as outliers independently for each clinical parameter and treated as missing values prior to regression analysis.

The clinical tests used in this study are summarized in Table S1. Sex, age, and a history of 36 specific diseases (Table S2) were included as covariates in the regression models. For a subset of specific tests, history of alcohol consumption and smoking were also incorporated as additional covariates, following the previous study ^33^. Linear regression analyses were performed using the lm() function in R (version 4.5.1). To account for multiple testing, the Bonferroni correction was applied, and statistical significance was determined based on the adjusted p-values.

### Genome-wide selection scan

SNPs associated with kidney function traits, including serum UA levels and eGFR, were obtained from PheWeb^52,53^. Variants showing genome-wide significance (*P* ≤ 5.0×10^−8^) were selected for further analyses. Genomic coordinates were converted from hg19 to hg38 using the UCSC liftOver tool to ensure compatibility with downstream functional annotation datasets^54^.

To prioritize renal trait-associated regions showing marked cross-population haplotype differentiation, we used previously reported XP-nSL scores between Jomon and CHB people^37,55^. Among genome-wide significant kidney trait-associated variants identified from PheWeb, we selected candidate regions containing at least 10 consecutive associated variants with an absolute XP-nSL score greater than 2.0. Within these candidate regions, we further calculated *f_2_* statistics to quantify allele-frequency differentiation between Jomon and CHB and prioritized variants with high *f_2_* ranks for subsequent annotation.

### *In silico* functional annotation and eQTL analysis

The potential regulatory roles of the identified candidate variants were assessed using multiple bioinformatics resources. eQTL (expression quantitative trait locus) data were retrieved from the GTEx database (V10)^56^ to evaluate the impact of each variant on gene expression across various human tissues. Through these platforms, we obtained information regarding chromatin accessibility (as determined by ATAC-seq or DNase-seq peaks) and histone modification marks across diverse tissue types through RegulomeDB (v2.2)^57^. These databases were also used to identify transcription factor binding motifs and genomic footprinting data within the regions encompassing the target SNPs.

### RNA-seq

PBMC samples separated by density gradient centrifugation were obtained from ToMMo. Total RNA was extracted from PBMCs using Maxwell RSC simplyRNA Cells Kit (Promega, Cat#AS1390) on Maxwell RSC Instrument (Promega, Cat#AS4500). The library prepared with Illumina Stranded mRNA Prep, Ligation (Illumina, Cat#20040534) and Illumina RNA UD Indexes Set B, Ligation (Illumina, Cat#20091657) was sequenced by a NovaSeq6000 System (Illumina). Sequencing was performed in a 2×100-bp PE configuration with a data output of about 40 million paired reads per sample. Library preparation and sequencing were performed in the National Institute of Genetics.

FASTQ files were preprocessed and subjected to quality control using fastp (v0.20.0) with adapter trimming and removal of the first base from both reads (-f 1 -F 1 -x)^58^. The duplication rate estimated using fastp ranged from 27.3% to 40.2%, and over 97.8% of the reads were successfully mapped to the reference genome (Table S10). Standard alignment and quality control metrics were assessed; however, no samples were excluded based on these metrics. The trimmed reads were aligned to the reference genome (GRCh38, Ensembl release 109 annotation) using HISAT2 (v2.2.1) with the --rna-strandness RF option^59^. Aligned reads were sorted by genomic coordinates using samtools 1.10^60^, and only properly paired reads were retained for downstream analyses. Read counts for genes were obtained from the aligned BAM files using featureCounts from the Subread package (v2.0.8) in paired-end mode (-p), requiring both ends to be mapped (-B), counting reads overlapping exons (-t exon)^61^. Genes with low expression levels were filtered using the filterByExpr function in the edgeR (4.8.0), taking into account the experimental group information, reducing the dataset from 62,710 to 17,190 genes^62,63^. Differential expression analysis was performed using DESeq2 (1.50.1) with a design formula including only the group variable (∼ group), comparing the upper-JAS group and the lower-JAS group^64^. Log2 fold changes for the lower-JAS group vs the upper-JAS group contrast were estimated and shrunk using the apeglm method^64,65^. Genes with |log2 fold change| ≥ 1 (fold change ≥ 2) and an adjusted p-value < 0.1 were considered significantly up- or down-regulated. These DEG entries were annotated with HGNC gene symbols while retaining Ensembl gene entries sharing the same gene symbol, resulting in a final set of 14,562 gene entries for functional enrichment analysis in Metascape using default parameters (GO Biological Processes), with all expressed genes possessing valid HGNC symbols used as the background set^66^.

RNA-seq count data generated by featureCounts were used to estimate the relative proportions of immune cell types in PBMCs to assess potential differences between sample groups. Gene length information was retrieved from biomaRt (Ensembl GRCh38, release 112) using the longest transcript per gene, and counts were converted to transcripts per million (TPM). Immune cell fractions were inferred using CIBERSORT via the immunedeconv R package with the LM22 signature matrix^41,67^, representing 22 human immune cell types. Genes without an HGNC gene symbol were excluded, and one entry per HGNC gene symbol was retained for CIBERSORT input. Comparisons of cell type fractions between the upper-JAS group (n = 5) and the lower-JAS group (n = 5) were performed using the Wilcoxon rank-sum test (exact = FALSE). Figures were generated using ggplot2 on data reshaped to long format for visualization.

### Establishment and culture of hiPSCs

The hiPSCs were generated from PBMCs, using SRV iPSC-4 Vector (TOKIWA-Bio, Cat#S1011696A) carrying *OCT4*, *SOX2, KLF4*, *MYC*, *NANOG, LIN28*, *EGFP,* and puromycin resistance gene. PBMCs were plated at 5.0x10^5^ cells into 96-well culture plates (Sumitomo Bakelite, Cat#MS-8096F) and pre-cultured in StemFit AK03N without C medium (Ajinomoto, Cat#AK03N) supplemented with Recombinant human SCF (50 ng/mL, Fuji film wako pure chemical corporation, Cat#197-15511), Recombinant human TPO (10 ng/mL, Fuji film wako pure chemical corporation, Cat#207-17581), Recombinant human Flt3L (20 ng/mL, Fuji film wako pure chemical corporation, Cat#061-05391), Recombinant human IL-6 (50 ng/mL, Fuji film wako pure chemical corporation, Cat#098-06041), Recombinant human IL-3 (20 ng/mL, Fuji film wako pure chemical corporation, Cat#090-05761), and Recombinant human G-CSF (10 ng/mL, Fuji film wako pure chemical corporation, Cat#072-06101) for 5 days. After pre-culture, 5x10^4^ cells were infected with this SRV iPSC-4 Vector at 3 multiplicity of infection (MOI), and plated into 6-well culture plates (Sumitomo Bakelite, Cat#MS-80060) with StemFit AK03N without C medium supplemented with 50 ng/mL Recombinant human SCF, 10 ng/mL Recombinant human TPO, 20 ng/mL Recombinant human Flt3L, 50 ng/mL Recombinant human IL-6, 20 ng/mL Recombinant human IL-3, 10 ng/mL Recombinant human G-CSF, and iMatrix-511 (0.5 μg/cm^2^, Matrixome, Cat#892011). On day 1 and 3 after infection, StemFit AK03N was added to each well. On day 6 after infection, medium was changed to StemFit AK03N with 10 μM Y-27632 (Fuji film wako pure chemical corporation, Cat#036-24023). On day 8, 10, and 12 after infection, the medium was changed to StemFit AK03N. The hiPSCs at passage 2 were subjected to cell sorting, and only GFP-negative cells were selected and passaged. hiPSCs were dissociated with TrypLE Express (Thermo Fisher Scientific, Cat#12604021) and seeded at 1.3 x 10^4^ cells/well in StemFit AK02N medium (Ajinomoto, Cat#AK02N) supplemented with 10 μM Y-27632 on 6-well iMatrix-511-coated culture plates. The next day, the medium was replaced with fresh StemFit AK02N medium. The established hiPSCs were maintained in StemFit AK02N medium on iMatrix-511-coated culture plates. All the cell culture was performed at 37°C, 5% CO2 in this study.

### SeV detection

Total RNA was extracted from PBMCs using Maxwell RSC simplyRNA Cells Kit on Maxwell RSC Instrument. SeV detection was performed with TaqMan Fast Virus 1-Step Master Mix (Thermo Fisher Scientific, Cat#4444427) on an ELITe InGenius system (ELITechGroup). For additional SeV detection, total RNA was extracted with RNeasy Mini Kit (Qiagen, Cat#74104) and reverse-transcribed into cDNA with the SuperScript IV VILO Master Mix with ezDNase Enzyme (invitrogen, Cat#11766050). Quantitative RT-PCR analyses were performed with SYBR Green qPCR Master Mix (Thermo Fisher Scientific, Cat#A66732) on a StepOne Real-Time PCR System (Applied Biosystems). Primer sequences are provided in Table S22^68-76^.

### Differentiation of hiPSCs into three germ layers

Differentiation of three germ layers was performed using the STEMdiff Trilineage Differentiation Kit (STEMCELL Technologies, Cat#ST-05230). hiPSCs were seeded at a density of 4.4 x 10^4^ cells/cm^2^ for endoderm and ectoderm differentiation, and 2.2 x 10^4^ cells/cm^2^ for mesoderm differentiation in StemFit AK03N medium supplemented with 10 μM Y-27632 and Matrigel (Corning, Cat#354277). The next day, as day 1 of differentiation, the culture medium was changed to differentiation medium. For endoderm and mesoderm differentiation, the medium was changed on day 1, 2, 3, and the cells were collected for immunocytochemistry on day 5. For ectoderm differentiation, the medium was changed on day 1, 2, 3, 5 and the cells were collected for immunocytochemistry on day 7.

### Characterization of hiPSCs

For immunocytochemistry, hiPSCs (passage number 5) and three germ layers were fixed with 4% paraformaldehyde (PFA) (Fuji film wako pure chemical corporation, Cat#163-20145), permeabilized with 0.1% Triton X-100 (Roche, Cat#10789704001), and blocked with 5% Normal Donkey Serum (Abcam, Cat#ab7475). For imaging, the cells were then incubated overnight with primary antibodies: anti-OCT4 (Abcam, Cat#ab19857, 1:150) and anti-SSEA4 (Abcam, Cat#ab16287, 1:375) at 4°C. For flow cytometry, the cells were then incubated with primary antibodies: OCT4 (Abcam, Cat#ab19857, 1:100), TRA-1-60 (Abcam, Cat#ab16288, 1:125), Pax6 (D3A9V) Rabbit Monoclonal Antibody (Cell Signaling Technology, Cat#60433, 1:50), Anti-Human Nestin Antibody (STEMCELL Technologies, Cat#60091, 1:50), TBXT antibody (Proteintech, Cat#20741-1-AP, 1:50), SOX17 antibody (Proteintech, Cat#24903-1-AP, 1:50), and Human CXCR4 Antibody (Bio-Techne, Cat#MAB170, 1:50) at room temperature. After washing with PBS, the cells were incubated with secondary antibodies: Donkey anti-Mouse IgG (H+L) Highly Cross-Adsorbed Secondary Antibody Alexa Fluor Plus 555 (Thermo Fisher Scientific, Cat#A32773, 1:500–1:1000), Donkey anti-Rabbit IgG (H+L) Highly Cross-Adsorbed Secondary Antibody Alexa Fluor Plus 647 (Thermo Fisher Scientific, Cat#A32795, 1:500–1:1000) at room temperature for 1 hour. Nuclei were counterstained with DAPI (Fuji film wako pure chemical corporation, Cat#340-07971). hiPSCs incubated with secondary antibodies alone, without primary antibodies, were used as a negative control. Images were acquired using a BZ-X700 (Keyence). Flow cytometry was performed with MA900 Multi-Application Cell Sorter (Sony).

### Differentiation of hiPSCs into cardiomyocytes

Differentiation of cardiomyocytes was performed using the PSC Cardiomyocyte Differentiation Kit (Thermo Fisher Scientific, Cat#A2921201). hiPSCs were seeded at a density of 0.5–2 x 10^4^ cells/cm^2^ in StemFit AK03N medium supplemented with 10 μM Y-27632 and iMatrix-511. The next day, as day 1 of differentiation, the culture medium was changed to StemFit AK03N without Y-27632. For differentiation, the medium was changed to Medium A on day 3, Medium B on day 5, and Maintenance Medium on day 7, 9, 11, and 13. The cells were collected for immunocytochemistry on day 15. For immunocytochemistry, cells were fixed with 4% PFA, permeabilized with 0.1% Triton X-100, and blocked with 5% Normal Donkey Serum. For imaging, the cells were then incubated overnight with primary antibodies: anti-Troponin T (Thermo Fisher Scientific, Cat#MA5-12960, 1:500) at 4°C. After washing with PBS, the cells were incubated with secondary antibodies: Donkey anti-Mouse IgG (H+L) Highly Cross-Adsorbed Secondary Antibody, Alexa Fluor Plus 555 (Thermo Fisher Scientific, Cat#A32773, 1:1000). Nuclei were counterstained with DAPI. Images were acquired using a CellVoyager CQ1 (Yokogawa Electric).

### Differentiation of hiPSCs into hepatocyte-like cells (HLCs) and validation of marker expression

Differentiation of hiPSCs into hepatocyte-like cells was performed following the protocol with some modifications^76,77^. Briefly, cells seeded at a density of 4.2 x 10^4^ cells/cm^2^ in iMatrix-511-coated 12- or 24-well culture plates were induced toward definitive endoderm (DE) for 2 days in RPMI 1640 (Nacalai Tesque, Cat#30264-56) supplemented with 1× B27 (Thermo Fisher Scientific, Cat#17504044), 100 ng/mL Activin A (Apro Science, Cat#GF-001-050L), 2 μM CHIR99021 (Cayman Chemical, Cat#13122), and 0.5 μM Sodium Butyrate (Nacalai Tesque, Cat# 12328-44), and then for 2 days in RPMI 1640 supplemented with 1× B27, 100 ng/mL Activin A, and 2 μM CHIR99021. For hepatic specification, DEs were cultured for 8 days in CTS KnockOut DMEM/F-12 (Thermo Fisher Scientific, Cat#A1370801) supplemented with 10% KnockOut Serum Replacement (Thermo Fisher Scientific, Cat#10828010), 0.5% GlutaMAX supplement (Thermo Fisher Scientific, Cat#35050-061), 0.5% MEM Non-Essential Amino Acids Solution (Fuji film wako pure chemical corporation, Cat#139-15651), 100 ng/mL Recombinant Human HGF (PeproTech, Cat#100-39), 1% DMSO (Nacalai Tesque, Cat#09659-14), and further cultured for 3 days in the presence of 100 nM Dexamethasone (Sigma-Aldrich, Cat#D4902). For hepatic maturation, cells were covered with a Matrigel (Corning, Cat#356237) overlay and maintained in Hepatocyte Culture Medium (HCM; Lonza, Cat#CC-3198), from which human EGF was omitted, supplemented with 20 ng/mL Oncostatin M (PeproTech, Cat#300-10).

For immunocytochemistry, cells were fixed with 4% PFA, permeabilized with 0.125% Triton X-100 (Sigma-Aldrich, Cat#X100), and blocked with 20% ImmunoBlock (KAC, Cat#KN001). For imaging, the cells were then incubated overnight with primary antibodies: Anti-albumin Antibody (Bethyl Laboratories, Cat#A80-229A, 1:150) and HNF-4α Antibody (Santa Cruz Biotechnology, Cat#sc-374229, 1:50) at 4. After washing with PBS, the cells were incubated with secondary antibodies: Donkey Anti-Goat IgG (H+L) Cross-Adsorbed, Alexa Fluor 488 (Thermo Fisher Scientific, Cat#A11055, 1:500) and Donkey anti-Mouse IgG (H+L) Highly Cross-Adsorbed Secondary Antibody, Alexa Fluor 488 (Thermo Fisher Scientific, Cat#A21202, 1:500). Nuclei were counterstained with DAPI (Dojindo, Cat#D523). Images were acquired using a BZ-X700 (Keyence).

Quantitative analysis of immunocytochemistry images was performed using ImageJ (Fiji) software (v2.1.0)^78^. DAPI images were converted to 8-bit format. Automated thresholding was applied using the Li algorithm, followed by binary conversion and watershed segmentation to separate clustered nuclei. The Analyze Particles function (size: 50– infinity pixel²) was used to define each nucleus as a Region of Interest (ROI) and recorded in the ROI Manager. The HNF4α images (16-bit) were overlaid with the DAPI-derived ROIs, and the Mean Gray Value was measured for each ROI. Background fluorescence was measured in cell-free regions of each image, and a threshold for HNF4α positivity was defined based on the background fluorescence. Nuclei with a Mean Gray Value above this threshold were classified as HNF4α-positive. Finally, the differentiation efficiency was calculated by dividing the number of HNF4α-positive nuclei by the total number of DAPI-positive nuclei for each image. One field of view per cell line was analyzed.

Total RNA was extracted from one to two wells per hiPSC or HLC line using RNeasy Mini Kit and reverse-transcribed into cDNA with the SuperScript IV VILO Master Mix with ezDNase Enzyme. Quantitative RT-PCR analyses were performed with SYBR Green qPCR Master Mix on a StepOne Real-Time PCR System (Applied Biosystems). Quantitative RT-PCR was performed in technical duplicate, and each independent hiPSC line was treated as one biological replicate (n = 5 per group). Gene expression levels were calculated using the 2^−ΔΔCT^ method with β*-ACTIN* as the internal control gene. Expression levels were normalized to J1 hiPSCs, which were set to 1, and are presented as log2 fold changes. Primer sequences are provided in Table S22.

### DNA extraction and Short Tandem Repeats (STR) analysis

Genomic DNA was extracted from PBMCs using TE Buffer Solution (Nacalai Tesque, Cat#06890-25) and Proteinase K (Nacalai Tesque, Cat#15679-06). Genomic DNA was extracted from hiPSCs using QIAamp DNA Mini Kit (Qiagen, Cat# 51304). To confirm the genetic identity between the donor PBMCs and the derived hiPSC lines, STR analysis was performed using the GenePrint 10 System (Promega, Cat#B9510). The analysis was conducted by BEX Co., Ltd. (Tokyo, Japan).

### Karyotype analysis

Metaphase spreads of the hiPSC lines were generated following established protocols ^79,80^. Briefly, hiPSC cultures were incubated with 0.05 μg/mL KaryoMAX Colcemid Solution (Thermo Fisher Scientific, Cat#15210-040) for 1.5 h to arrest cells in metaphase. Cells were subsequently collected and subjected to hypotonic treatment with 0.075 M KCl containing 0.1% (w/v) trisodium citrate at 37°C for 22 min, then fixed using Carnoy’s solution. The cell suspensions were dropped onto glass slides. Chromosomal analysis, including determination of chromosome number and detection of structural abnormalities, was performed by QFH-banding or mFISH, depending on the hiPSC line. QFH-banding was conducted using Quinacrine mustard dihydrochloride Fluorescent Dye (Sigma-Aldrich, Cat#Q2876) and Hoechst 33258 solution Fluorescent DNA dye (Sigma-Aldrich, Cat# 94403)^81^. For mFISH analysis, metaphase spreads were hybridized with the 24XCyte human mFISH probe set (MetaSystems) according to the manufacturer’s instructions^82,83^. For each hiPSC line, more than 10 metaphase spreads were imaged and analyzed using either a Leica DM5000 B fluorescence microscope equipped with a CCD camera and CW4000 software (Leica Microsystems) or a fluorescent microscope equipped with a CCD camera and ISIS image analysis systems (Carl Zeiss microscopy).

### Mycoplasma tests

The hiPSC culture medium at passage 8 was tested for mycoplasma contamination by CycleavePCR Mycoplasma Detection Kit (Takara Bio, Cat#CY232). Detection was performed on the StepOnePlus Real-Time PCR System (Applied Biosystems). All hiPSC lines used in this study were confirmed to be negative for mycoplasma.

### Triglyceride measurement

Triglyceride (TG) in HLC was measured by EnzyChrom Triglyceride Assay Kit (BioAssay Systems, Cat#ETGA-200). Cells from four wells of a 12-well plate were pooled and lysed with 5% Triton X-100 (Sigma-Aldrich, Cat#X100). To normalize TG content to total cellular protein, protein concentrations were determined using TaKaRa BCA Protein Assay Kit (Takara Bio, Cat#T9300A), and equal amounts of total protein were used for the assay. OD was measured by GloMax Discover Microplate Reader (Promega, Cat#GM3000).

## Resource availability

### Lead contact

Further information and requests for resources and reagents should be directed to and will be fulfilled by the Lead Contact, Hiroki Oota.

### Materials availability

The hiPSCs generated in this study are available upon reasonable request.

### Data and code availability

The RNA-seq data generated in this study are being deposited in the NBDC Human Database. The accession number will be provided once registration is complete. The genomic data of the ToMMo were used under license for the current study and are not publicly available. Access to these data can be requested through the ToMMo data distribution portal. Researchers who obtain approval from ToMMo to access the original genomic and clinical datasets can identify the specific samples used in this study by recalculating the JAS according to the methods described herein. The analysis scripts used for processing the data (e.g., DESeq2 pipeline) are provided in the Supplemental Information. Any additional information required to reanalyze the data reported in this article is available from the lead contact upon request.

## Acknowledgments

This study was supported by JSPS KAKENHI Grant-in-Aid for Scientific Research (A) to 21H04779 and 25H00999 to H.O., JSPS KAKENHI Grant Number JP22H04925 (PAGS) to H.O., The Mitsubishi Foundation to H.O., JST SPRING Grant Number JPMJSP2108 to Y.N., Rikaken Holdings Young Researcher Support Grant to Y.N., JSPS KAKENHI Grant Number 26KJ0827 to Y.N. Computations were partially performed on the NIG supercomputer at ROIS National Institute of Genetics. This research was supported (in part) by the Japan Agency for Medical Research and Development, AMED under Grant Number JP21tm0424601. We thank the Uehiro Future Scientists Program at the CiRA Foundation for supporting the establishment of the iPSC lines.

## Author contributions

Y.N., Y.W., and H.O. conceived the study. Y.N. and Y.W. designed and performed data analyses. Y.N. conducted iPSC-based experiments and curated the data. A.T. performed sequencing. H.T. performed karyotype analysis. Y.N. wrote the original draft of the manuscript with support from Y.W. and H.O. A.H., T.K., K.H., K. KU., T.T., and J.O. provided and developed the methodology and reviewed and edited the manuscript. M.T., Y.H., M.I., K.KO., and T.I. contributed to discussions and reviewed and edited the manuscript. H.O. supervised the project. All authors have read and approved the final article.

## Declaration of generative AI and AI-assisted technologies in the writing process

Gemini and ChatGPT were used to improve the language quality and readability. During the preparation of this manuscript, the authors reviewed and refined the content and take full responsibility for the content of the publication.

## Figure legends

Figure S1: (A) Distribution of all clinical values. Data points exceeding four standard deviations (4SD) from the mean were excluded prior to plotting. (B) Association between JAS and all clinical parameters analyzed in this study. Linear regression analyses were performed between JAS and 31 clinical parameters. The upward- and downward-pointing triangles indicate positive and negative associations, respectively. The y-axis shows the statistical significance of each association (−log10 p-value). (C) Associations between JAS and clinical test values. Scatter plots show the relationship between JAS and each clinical test value, with each point representing an individual.

Figure S2: Estimated proportions of immune cell subtypes in PBMCs. Estimated proportions of 22 immune cell types inferred using CIBERSORT. Each dot represents an individual donor (n = 5 per group). Group differences were assessed using the Wilcoxon rank-sum test.

Figure S3: Flow cytometric analysis of pluripotency markers (OCT4 and TRA-1-60) in 10 hiPSC lines. A hiPSC line (T1) incubated with secondary antibodies alone was used as a negative control.

Figure S4: Flow cytometric analysis of ectodermal (PAX6 and Nestin) markers in 10 hiPSC lines differentiated into the ectoderm linage. A hiPSC line (T1) incubated with secondary antibodies alone was used as a negative control.

Figure S5: Flow cytometric analysis of mesodermal (TBXT (Brachyury) and CXCR4) markers in 10 hiPSC lines differentiated into the mesoderm linage. A hiPSC line (T1) incubated with secondary antibodies alone was used as a negative control.

Figure S6: Flow cytometric analysis of endodermal markers (SOX17 and CXCR4) in 10 hiPSC lines differentiated into the endoderm linage. A hiPSC line (T1) incubated with secondary antibodies alone was used as a negative control.

Figure S7: Immunofluorescence images of the cardiomyocyte marker (Troponin T) in 10 hiPSC lines differentiated into cardiomyocyte-like cells. Scale bar: 300 μm.

Figure S8: QFH-banding and mFISH of chromosomes obtained from metaphase spreads of 10 hiPSC lines.

Figure S9: Immunofluorescence images of the hepatic markers (albumin (ALB) and HNF4α) in 10 hiPSC lines differentiated into hepatocyte-like cells. Scale bars: 200 μm.

Figure S10: Intracellular triglyceride content in hiPSCs derived from individuals with the upper- and lower-JAS groups. No significant difference was observed between the upper- and lower-JAS groups by the Mann–Whitney U test (n = 5 independent hiPSC lines per group, with four wells pooled for each line).

