## Supplementary figures and images for "An iPSC-based experimental framework for investigating phenotypic variation associated with Jomon ancestry in the present-day Japanese population"

### Supplementary Figures

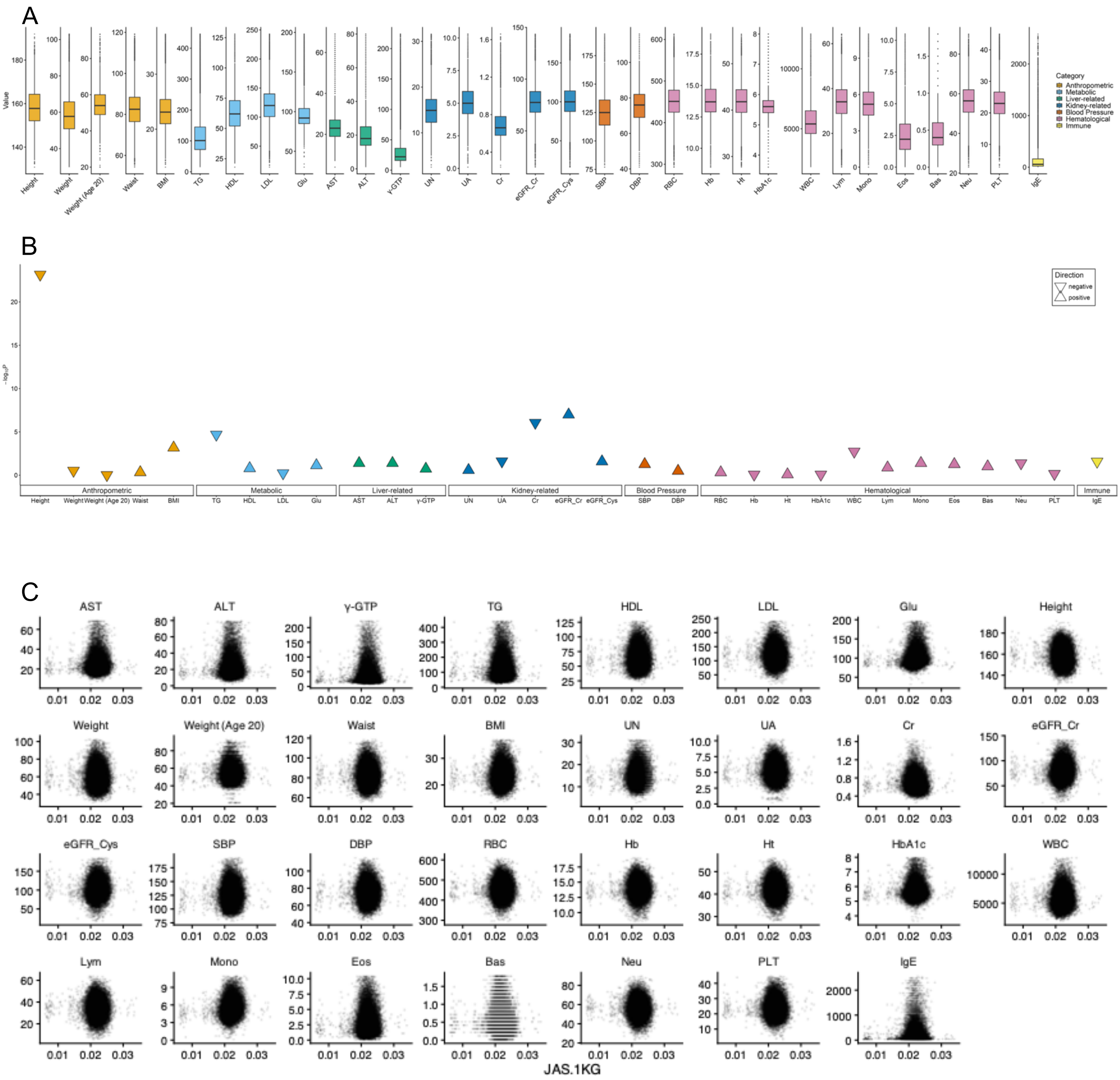

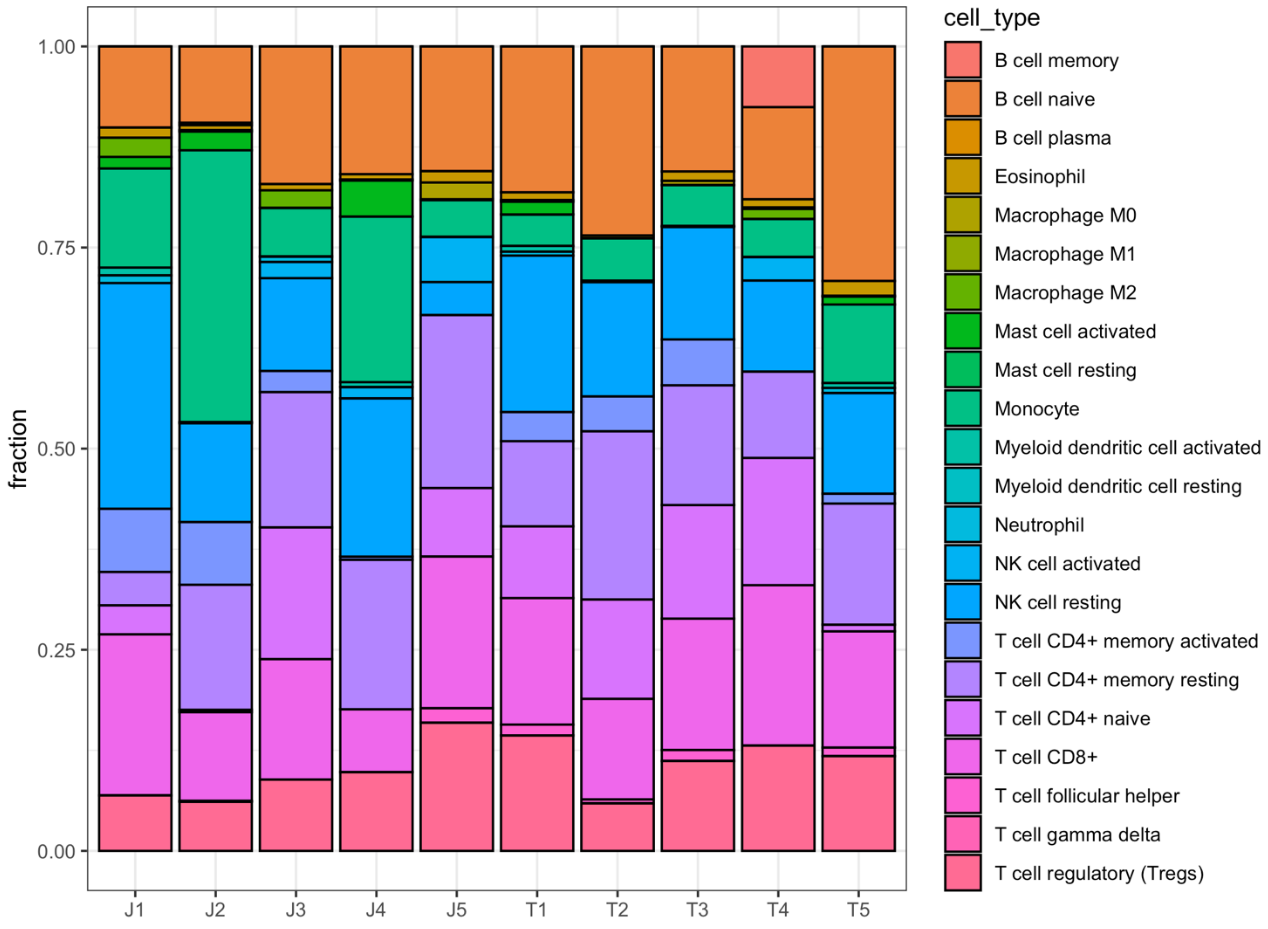

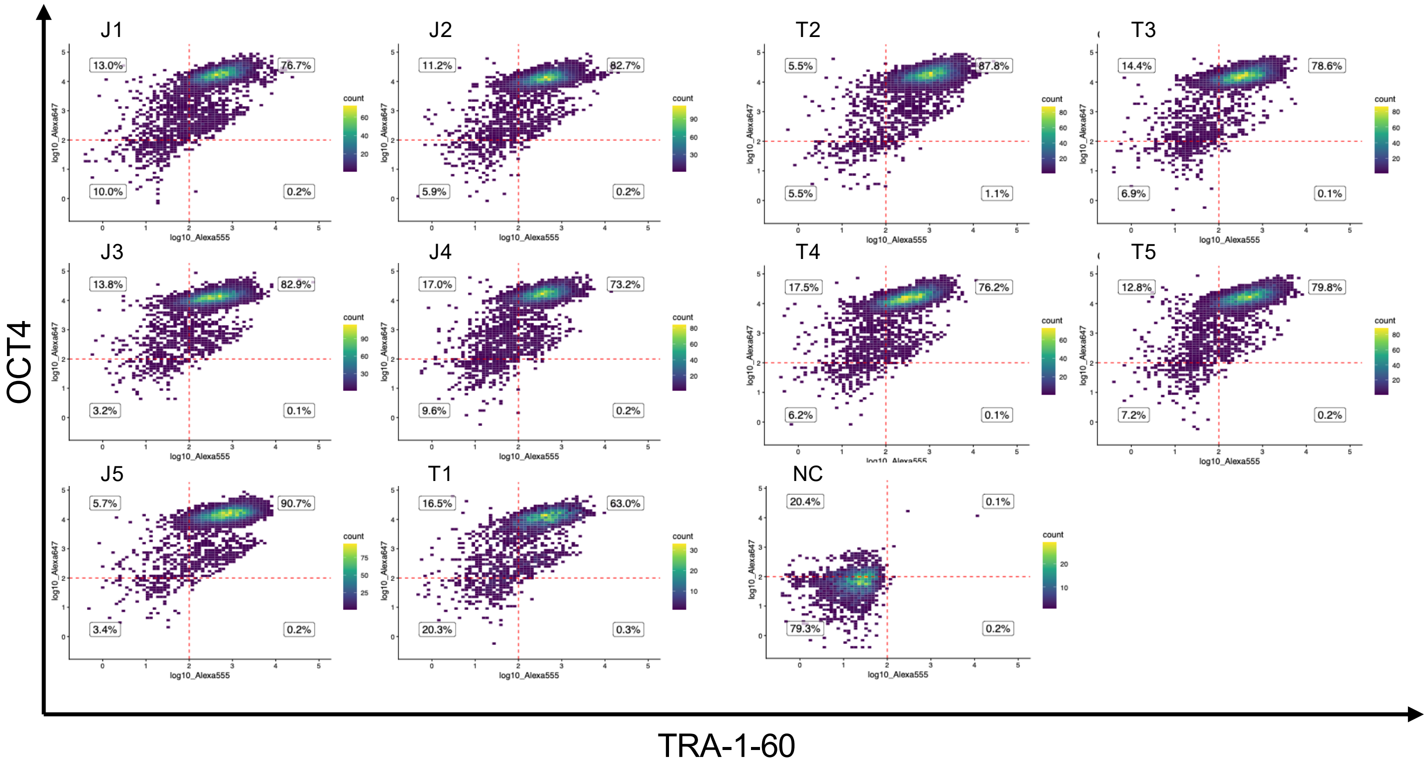

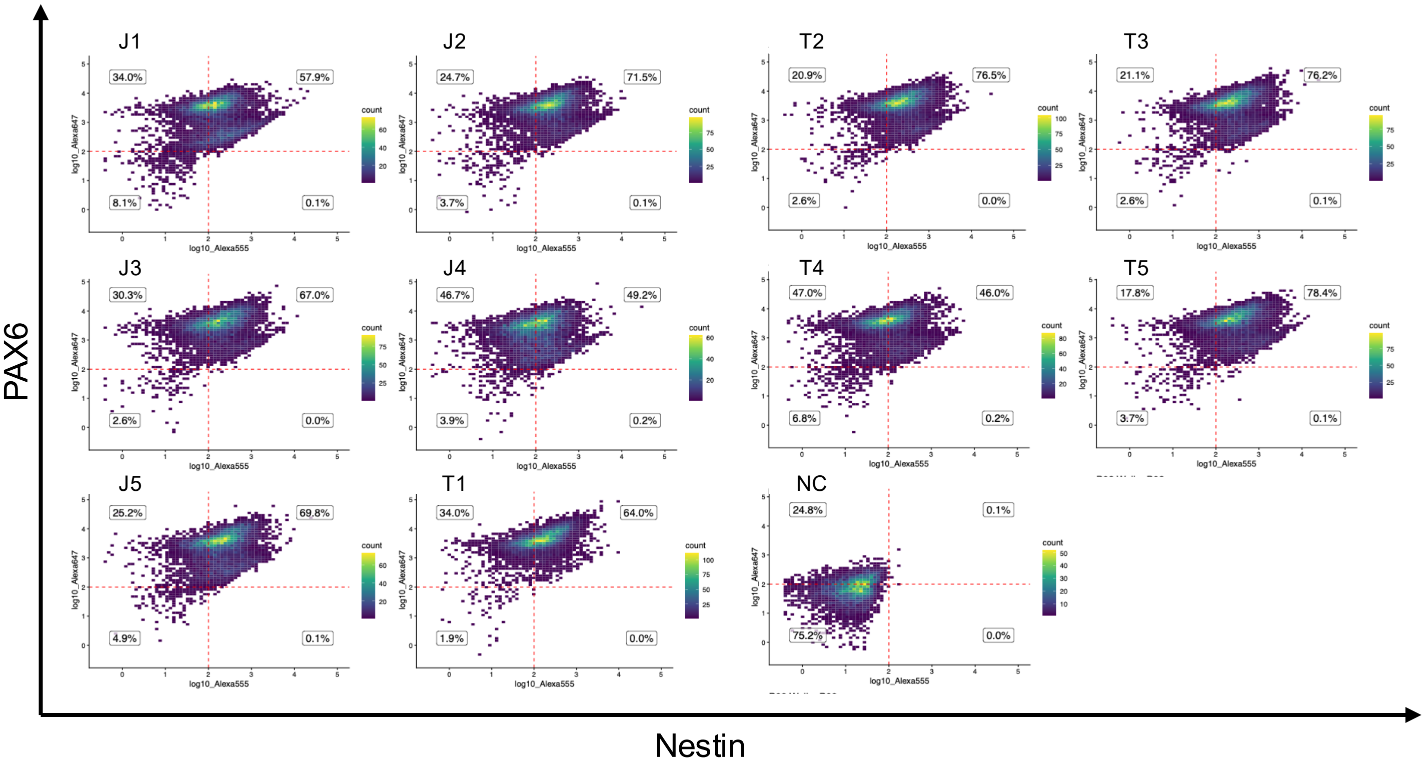

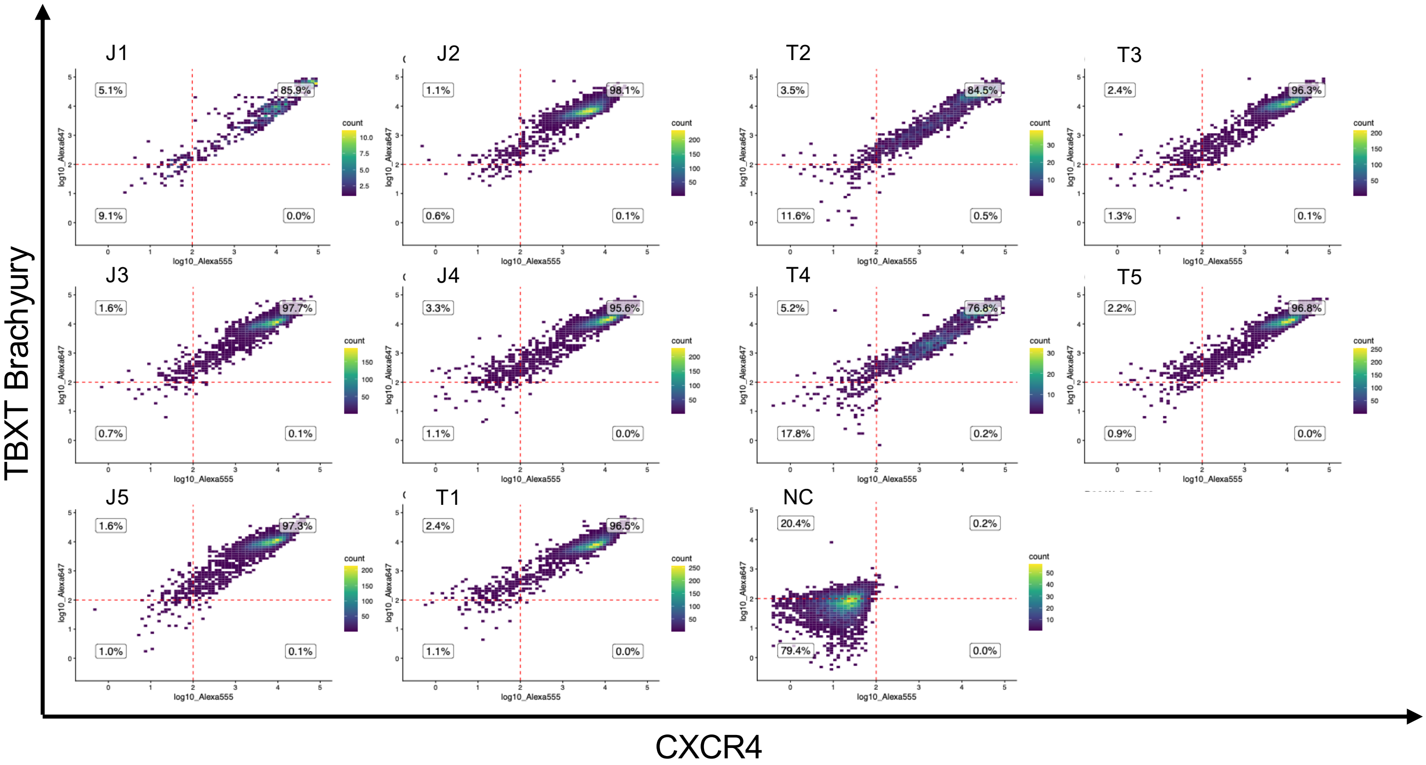

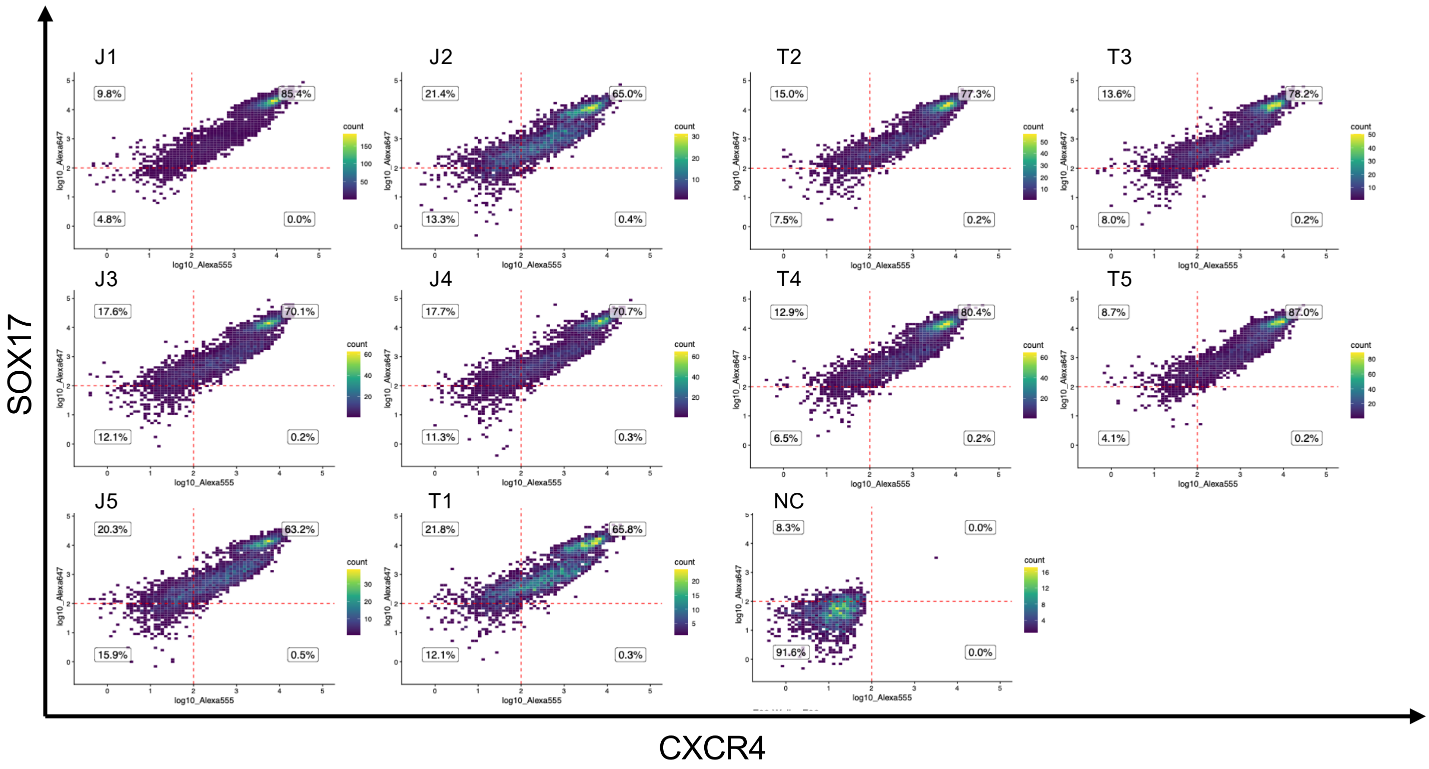

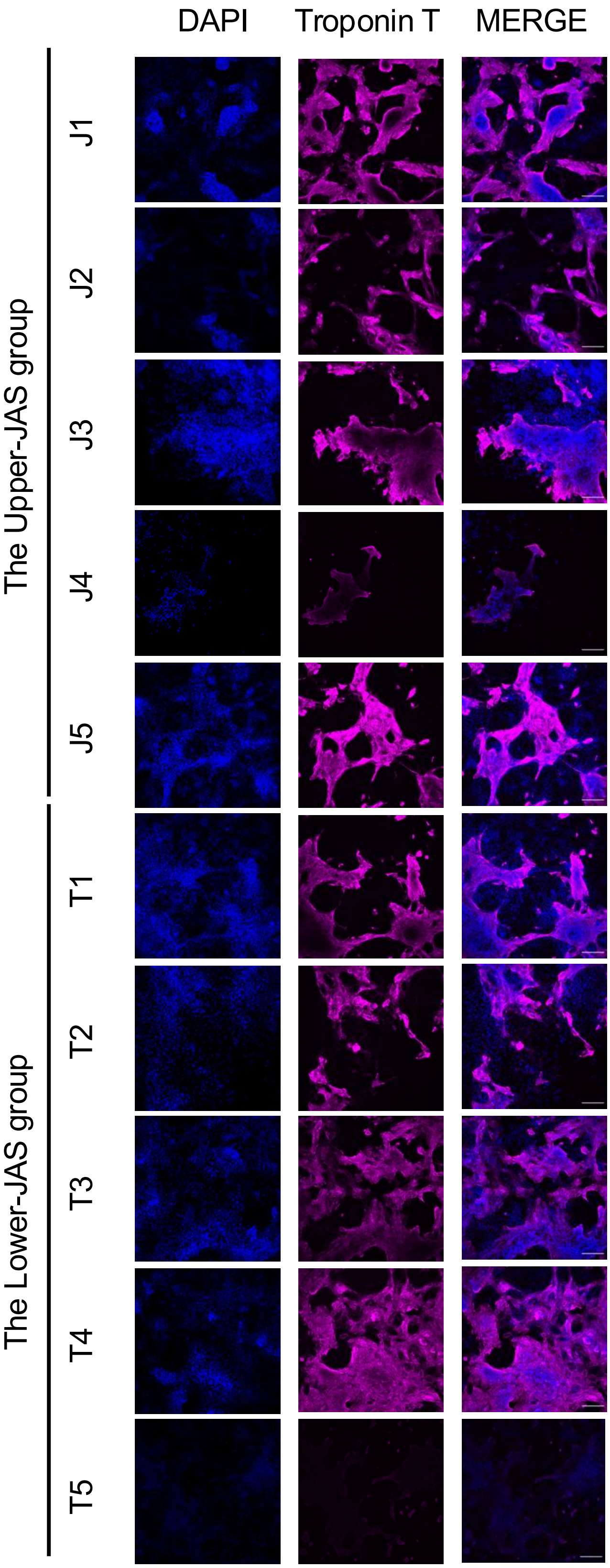

Figure S8

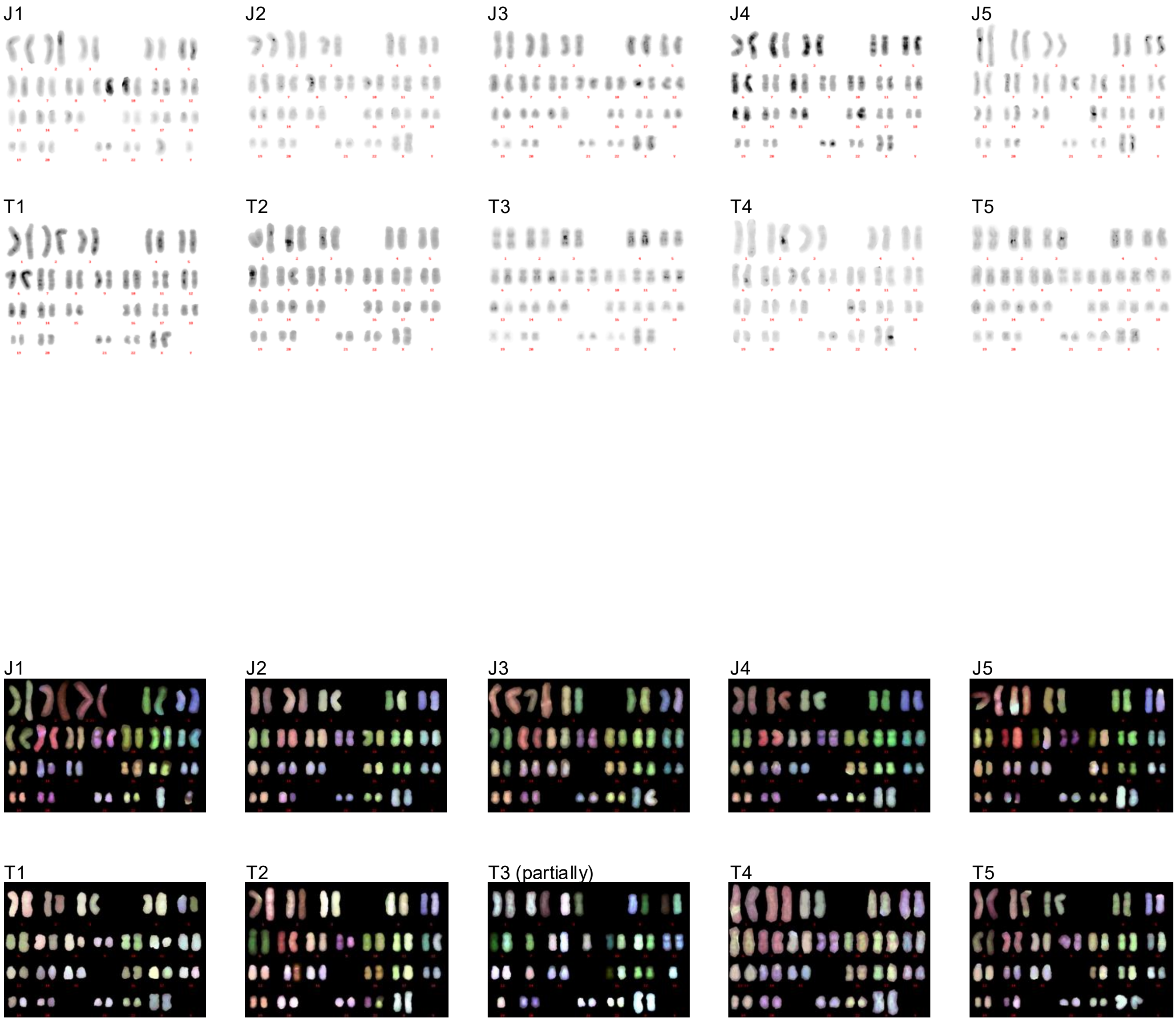

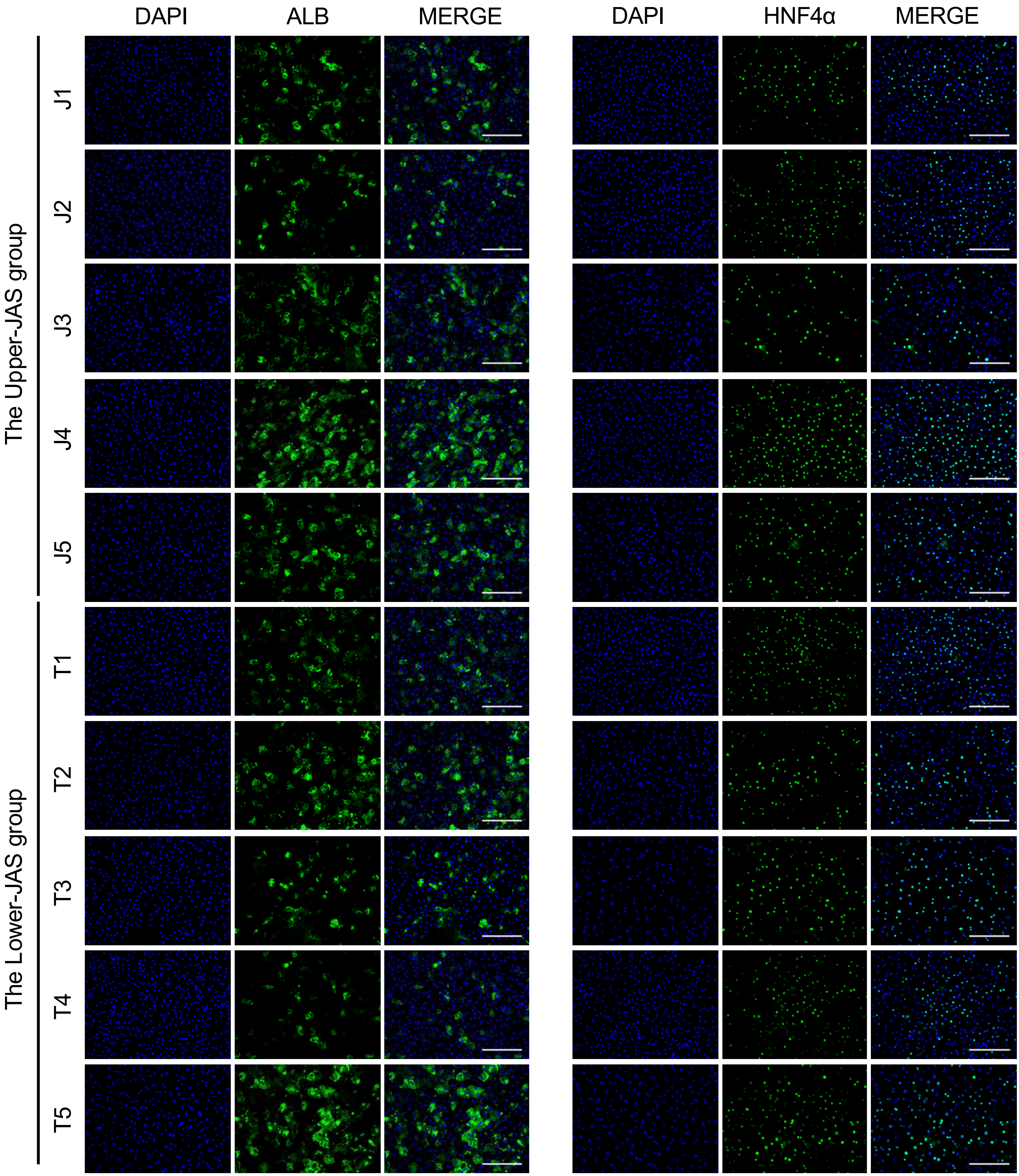

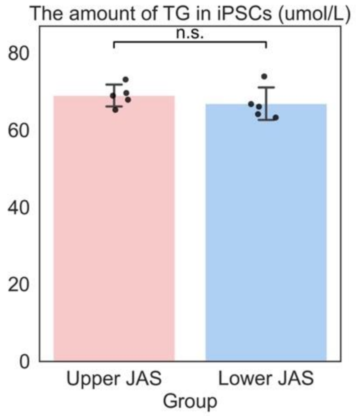
