## Supplementary Information for "An iPSC-based experimental framework for investigating phenotypic variation associated with Jomon ancestry in the present-day Japanese population"

Supplemental Information

Data S1. Data preprocessing and linear regression analysis of clinical phenotypes associated with Jomon ancestry.

import pandas as pd

#====================

\# Define the analysis population

\# ====================

df1 = pd.read_csv("laboratory_test.csv", low_memory=False)

df2 = pd.read_csv("health_checkup.csv", low_memory=False)

df3 = pd.read_csv("lifestyle_questionnaire.csv", low_memory=False)

df4 = pd.read_csv("JAS_PCA.csv", low_memory=False)

\# Remove unnecessary columns

df1 = df1.drop(columns=["source_path", "reservation_number", "course_code"])

df2 = df2.drop(columns=["source_path", "reservation_number", "course_code"])

df3 = df3.drop(columns=["source_path", "reservation_number"])

\# Standardize questionnaire column names

new_cols = []

for c in df3.columns:

try:

new_cols.append(str(int(float(c))))

except:

new_cols.append(str(c))

df3.columns = new_cols

\# Select questionnaire variables

questionnaire_cols = ["ID", "405010", "405040", "405050", "405060", "405070", "405080", "405090", "405131", "406000", "406250", "410800", "410880", "412650", "412730", "412810", "412890", "412970", "413050", "413220", "413300", "414510", "415650", "415730", "415810", "415890", "415970", "416050", "416130", "416210", "416290", "416370", "416450", "416620", "416700", "416780", "416860", "416940", "417120", "417200", "417280", "417360", "417440", "417520", "417600", "417680", "417760", "417840"]

df3 = df3[questionnaire_cols]

\# Merge datasets by participant ID

merged = pd.merge(df1, df2, on="ID", how="outer")

merged = pd.merge(merged, df3, on="ID", how="outer")

merged = pd.merge(merged, df4, on="ID", how="outer")

\# Remove duplicated columns generated during merging

merged = merged.loc[:, ~merged.columns.str.endswith("_y")]

merged.columns = merged.columns.str.replace("_x", "", regex=False)

\# Remove columns containing no data

merged = merged.dropna(axis=1, how="all")

\# ====================

\# Restrict to the analysis population

\# ====================

\# Restrict to East Asian participants

non_east_asian_codes = [9643, 9608, 9076, 9152, 9360, 9585, 9586, 9600, 9764, 9804]

merged = merged[~merged["405131"].isin(non_east_asian_codes)]

\# Remove variables not used in the analysis

merged = merged.drop(columns=["405131", "405080", "405090", "240671.00", "240691.00", "416210"])

\# Remove related individuals

kinship_ids = pd.read_csv("king.kinship.maf0.1.out.id", header=None)[0]

merged = merged[~merged["ID"].isin(kinship_ids)]

\# Remove participants with missing values

merged = merged.dropna()

\# Remove participant identifiers

merged = merged.drop(columns=["ID", "date"])

\# Save analysis dataset

merged.to_csv("analysis_population.csv", index=False)

\# ====================

\# Linear regression analysis

\# ====================

library(car)

\# ============================================================

\# Load analysis dataset

\# ============================================================

df <- read.csv("analysis_population.csv", check.names = FALSE)

\# ============================================================

\# Define categorical covariates

\# ============================================================

categorical_vars <- c("course_code", "405010", "406000", "406250", "410800", "410880", "412650", "412730", "412810", "412890", "412970", "413050", "413220", "413300", "414510", "415650", "415730", "415810", "415890", "415970", "416050", "416130", "416290", "416370", "416450", "416620", "416700", "416780", "416860", "416940", "417120", "417200", "417280", "417360", "417440", "417520", "417600", "417680", "417760", "417840")

for (v in categorical_vars) {df[[v]] <- factor(df[[v]])}

\# ============================================================

\# Assess multicollinearity

\# ============================================================

vif_covariates <- c("course_code", "405010", "405040", "406000", "406250", "410800", "410880", "412650", "412730", "412810", "412890", "412970", "413050", "413220", "413300", "414510", "415650", "415730", "415810", "415890", "415970", "416050", "416130", "416290", "416370", "416450", "416620", "416700", "416780", "416860", "416940", "417120", "417200", "417280", "417360", "417440", "417520", "417600", "417680", "417760", "417840")

vif_formula <- paste("JAS.1KG ~", paste(vif_covariates, collapse = " + "))

vif_model <- lm(as.formula(vif_formula), data = df)

vif_result <- vif(vif_model)

write.csv(vif_result, "vif_results.csv")

\# ============================================================

\# Define covariates for regression models

\# ============================================================

covariates <- c("course_code", "405010", "405040", "410800", "410880", "412650", "412730", "412810", "412890", "412970", "413050", "413220", "413300", "414510", "415650", "415730", "415810", "415890", "415970", "416050", "416130", "416290", "416370", "416450", "416620", "416700", "416780", "416860", "416940", "417120", "417200", "417280", "417360", "417440", "417520", "417600", "417680", "417760", "417840")

\# ============================================================

\# Remove extreme values

\# ============================================================

outlier_variables <- c("101141.00", "101151.01", "240321.00", "240341.00", "240371.00", "240411.00", "241051.00", "241081.00", "241111.00", "241351.00", "241371.00", "241411.00", "241451.00", "242791.00", "405070", "240521.00", "240541.00", "240781.00", "240811.00", "240841.00", "240871.00", "240981.00", "242801.00", "242811.00", "242821.00", "242831.00", "242841.00", "242851.00", "241141.00", "241171.00", "241201.00")

for (v in outlier_variables) {

z_scores <- scale(df[[v]])

outlier_idx <- which(

abs(z_scores) > 4 &

!is.na(z_scores)

)

if (length(outlier_idx) > 0) {

df[outlier_idx, v] <- NA

}

}

\# ============================================================

\# Linear regression

\# ============================================================

clinical_traits <- c("101151.01", "240321.00", "240341.00", "240371.00", "240411.00", "241051.00", "241081.00", "241111.00", "241351.00", "241371.00", "241411.00", "241451.00", "242791.00", "405070")

for (trait in clinical_traits) {

formula <- paste(

"`", trait, "` ~ ",

paste(c("JAS.1KG", covariates), collapse = " + "),

sep = ""

)

model <- lm(

as.formula(formula),

data = df

)

write.csv(summary(model)$coefficients, paste0("model_", trait, ".csv"), row.names = TRUE)}

\# ============================================================

\# Linear regression with smoking as a covariate

\# ============================================================

smoking_traits <- c("101141.00", "240521.00", "240541.00", "240781.00", "240811.00", "240841.00", "240871.00", "240981.00", "242801.00", "242811.00", "242821.00", "242831.00", "242841.00", "242851.00")

for (trait in smoking_traits) {

formula <- paste(

"`", trait, "` ~ ",

paste(

c("JAS.1KG", covariates, "406250"),

collapse = " + "

),

sep = ""

)

model <- lm(

as.formula(formula),

data = df

)

write.csv(

summary(model)$coefficients,

paste0("model_smoking_", trait, ".csv"),

row.names = TRUE

)

}

\# ============================================================

\# Linear regression with alcohol consumption as a covariate

\# ============================================================

liver_traits <- c("241141.00", "241171.00", "241201.00")

for (trait in liver_traits) {

formula <- paste(

"`", trait, "` ~ ",

paste(

c("JAS.1KG", covariates, "406000"),

collapse = " + "

),

sep = ""

)

model <- lm(

as.formula(formula),

data = df

)

write.csv(summary(model)$coefficients, paste0("model_alcohol_", trait, ".csv"), row.names = TRUE)

}

Data S2. RNA-seq analysis scripts for preprocessing, read alignment, gene expression quantification, and differential expression analysis of PBMCs

#!/bin/bash

FASTQ_DIR="../fastq"

INDEX_PATH="../reference/grch38/genome"

GTF_PATH="../reference/Homo_sapiens.GRCh38.109.gtf"

SAMPLES=(J1 J2 J3 J4 J5 T1 T2 T3 T4 T5)

for SAMPLE in "${SAMPLES[@]}"; do

echo "Processing ${SAMPLE}..."

R1_FILES=$(find "${FASTQ_DIR}" -type f -name "${SAMPLE}_*_R1_001.fastq.gz" | sort)

R2_FILES=$(find "${FASTQ_DIR}" -type f -name "${SAMPLE}_*_R2_001.fastq.gz" | sort)

zcat ${R1_FILES} | gzip > "${SAMPLE}_R1.fastq.gz"

zcat ${R2_FILES} | gzip > "${SAMPLE}_R2.fastq.gz"

fastp -i "${SAMPLE}_R1.fastq.gz" -I "${SAMPLE}_R2.fastq.gz" -o "${SAMPLE}_R1.preprocessed.fastq.gz" -O "${SAMPLE}_R2.preprocessed.fastq.gz" -h "${SAMPLE}.fastp.html" -j "${SAMPLE}.fastp.json" -f 1 -F 1 -x

hisat2 --rna-strandness RF -x "${INDEX_PATH}" -1 "${SAMPLE}_R1.preprocessed.fastq.gz" -2 "${SAMPLE}_R2.preprocessed.fastq.gz" -S "${SAMPLE}.sam"

samtools sort -o "${SAMPLE}.sorted.bam" "${SAMPLE}.sam"

samtools view -f 0x2 -b "${SAMPLE}.sorted.bam" > "${SAMPLE}.paired.bam"

samtools index "${SAMPLE}.paired.bam"

samtools flagstat "${SAMPLE}.paired.bam" > "${SAMPLE}.flagstat.txt"

rm "${SAMPLE}.sam"

done

featureCounts -p --countReadPairs -B -T 4 -t exon -g gene_id -a "${GTF_PATH}" -o featureCounts.txt J1.paired.bam J2.paired.bam J3.paired.bam J4.paired.bam J5.paired.bam T1.paired.bam T2.paired.bam T3.paired.bam T4.paired.bam T5.paired.bam

\# Differential expression analysis of PBMC RNA-seq

library(edgeR)

library(DESeq2)

library(apeglm)

library(org.Hs.eg.db)

library(AnnotationDbi)

library(dplyr)

counts <- read.csv("featureCounts_repname.csv", row.names = 1, check.names = FALSE)

group <- factor(ifelse(grepl("^J", colnames(counts)), "upper_JAS", ifelse(grepl("^T", colnames(counts)), "lower_JAS", NA)), levels = c("upper_JAS", "lower_JAS"))

coldata <- data.frame(row.names = colnames(counts), group = group)

y <- DGEList(counts = counts, group = group)

keep <- filterByExpr(y, group = group)

counts_filtered <- counts[keep, ]

message("Number of genes before filtering: ", nrow(counts))

message("Number of genes after filtering: ", nrow(counts_filtered))

dds <- DESeqDataSetFromMatrix(countData = counts_filtered, colData = coldata, design = ~ group)

dds <- DESeq(dds)

res <- lfcShrink(dds, coef = "group_lower_JAS_vs_upper_JAS", type = "apeglm")

ids <- rownames(res)

ids_nover <- sub("[.].*", "", ids)

symbols <- mapIds(org.Hs.eg.db, keys = ids_nover, column = "SYMBOL", keytype = "ENSEMBL", multiVals = "first")

res_df <- as.data.frame(res)

res_df$ENSEMBL <- ids_nover

res_df$SYMBOL <- symbols

res_df <- res_df[!is.na(res_df$SYMBOL), ]

fdr_cutoff <- 0.1; log2fc_cutoff <- 1

up_genes <- res_df %>% filter(!is.na(padj), padj < fdr_cutoff, log2FoldChange >= log2fc_cutoff) %>% pull(SYMBOL) %>% unique()

down_genes <- res_df %>% filter(!is.na(padj), padj < fdr_cutoff, log2FoldChange <= -log2fc_cutoff) %>% pull(SYMBOL) %>% unique()

write.csv(res_df, "DEG_results.csv", row.names = FALSE)

write.csv(data.frame(Gene = up_genes), "upregulated_genes.csv", row.names = FALSE)

write.csv(data.frame(Gene = down_genes), "downregulated_genes.csv", row.names = FALSE)

vsd <- vst(dds, blind = FALSE)

all_genes <- rownames(vsd)

all_genes_nover <- sub("[.].*", "", all_genes)

gene_symbols <- mapIds(org.Hs.eg.db, keys = all_genes_nover, column = "SYMBOL", keytype = "ENSEMBL", multiVals = "first")

background_genes <- na.omit(data.frame(Ensembl = all_genes_nover, SYMBOL = gene_symbols))

write.csv(background_genes, "background_genes.csv", row.names = FALSE)

\# Immune cell deconvolution of PBMC RNA-seq

library(biomaRt)

library(immunedeconv)

library(dplyr)

library(tidyr)

library(ggplot2)

counts <- read.csv("featureCounts_repname.csv", row.names = 1, check.names = FALSE)

mart <- useEnsembl(biomart = "genes", dataset = "hsapiens_gene_ensembl", GRCh = 38)

gene_info <- getBM(attributes = c("ensembl_gene_id", "ensembl_transcript_id", "hgnc_symbol", "transcript_length"), mart = mart)

gene_lengths <- gene_info %>% filter(hgnc_symbol != "", !is.na(transcript_length)) %>% group_by(ensembl_gene_id) %>% slice_max(transcript_length, n = 1, with_ties = FALSE) %>% ungroup()

counts$ENSEMBL <- sub("[.].*", "", rownames(counts))

counts_annotated <- counts %>% left_join(gene_lengths, by = c("ENSEMBL" = "ensembl_gene_id")) %>% filter(!is.na(transcript_length), hgnc_symbol != "")

count_matrix <- counts_annotated %>% select(hgnc_symbol, all_of(setdiff(colnames(counts), "ENSEMBL"))) %>% group_by(hgnc_symbol) %>% summarise(across(everything(), sum), .groups = "drop")

rownames(count_matrix) <- count_matrix$hgnc_symbol

count_matrix <- count_matrix %>% select(-hgnc_symbol)

gene_length_matrix <- counts_annotated %>% select(hgnc_symbol, transcript_length) %>% distinct(hgnc_symbol, .keep_all = TRUE)

rownames(gene_length_matrix) <- gene_length_matrix$hgnc_symbol

gene_length_matrix <- gene_length_matrix[rownames(count_matrix), , drop = FALSE]

rpk <- sweep(as.matrix(count_matrix), 1, gene_length_matrix$transcript_length / 1000, "/")

tpm <- sweep(rpk, 2, colSums(rpk), "/") * 1e6

tpm_for_deconv <- as.data.frame(tpm)

tpm_for_deconv <- cbind(Gene = rownames(tpm_for_deconv), tpm_for_deconv)

rownames(tpm_for_deconv) <- NULL

immune_fractions <- deconvolute(tpm_for_deconv, method = "cibersort")

immune_fractions <- immune_fractions %>% column_to_rownames("cell_type")

sample_names <- colnames(tpm)

group <- ifelse(grepl("^J", sample_names), "upper_JAS", "lower_JAS")

results <- data.frame()

for (cell_type in rownames(immune_fractions)) {

values <- as.numeric(immune_fractions[cell_type, sample_names])

upper <- values[group == "upper_JAS"]

lower <- values[group == "lower_JAS"]

test <- wilcox.test(upper, lower, exact = FALSE)

results <- rbind(results, data.frame(cell_type = cell_type, p_value = test$p.value))

}

write.csv(immune_fractions, "CIBERSORT_cell_fractions.csv")

write.csv(results, "CIBERSORT_Wilcoxon_results.csv", row.names = FALSE)
